# Joint *ex vivo* MRI and histology detect iron-rich cortical gliosis in Tau and TDP-43 proteinopathies

**DOI:** 10.1101/2021.04.14.439064

**Authors:** M. Dylan Tisdall, Daniel T. Ohm, Rebecca Lobrovich, Sandhitsu R. Das, Gabor Mizsei, Karthik Prabhakaran, Ranjit Ittyerah, Sydney Lim, Corey T. McMillan, David A. Wolk, James Gee, John Q. Trojanowski, Edward B. Lee, John A. Detre, Paul Yushkevich, Murray Grossman, David J. Irwin

**Affiliations:** Radiology, Perelman School of Medicine, University of Pennsylvania; Neurology, Perelman School of Medicine, University of Pennsylvania; Pathology and Laboratory Medicine, Perelman School of Medicine, University of Pennsylvania

**Author notes:** Corresponding authors: David J. Irwin, Frontotemporal Degeneration Center (FTDC), University of Pennsylvania Perelman School of Medicine, Hospital of the University of Pennsylvania, 3600 Spruce Street, Philadelphia, PA 19104, M. Dylan Tisdall, Department of Radiology, University of Pennsylvania Perelman School of Medicine, D406 Richards Building, 3700 Hamilton Walk, Philadelphia, PA 19104.

**Keywords:** Frontotemporal lobar degeneration, Alzheimer’s disease, iron, *ex vivo* MRI, histopathology

## Abstract

Frontotemporal lobar degeneration (FTLD) is a heterogeneous spectrum of age-associated neurodegenerative diseases that include two main pathologic categories of tau (FTLD-Tau) and TDP-43 (FTLD-TDP) proteinopathies. These distinct proteinopathies are often clinically indistinguishable during life, posing a major obstacle for diagnosis and emerging therapeutic trials tailored to disease-specific mechanisms. Moreover, MRI-derived measures have had limited success to date discriminating between FTLD-Tau or FTLD-TDP. T2*-weighted (T2*w) *ex vivo* MRI has previously been shown to be sensitive to non-heme iron in healthy intracortical lamination and myelin, and to pathological iron deposits in amyloid-beta plaques and activated microglia in Alzheimer’s disease (AD). However, an integrated, *ex vivo* MRI and histopathology approach is understudied in FTLD. We apply joint, whole-hemisphere *ex vivo* MRI at 7T and histopathology to the study autopsy-confirmed FTLD-Tau (n=3) and FTLD-TDP (n=2), relative to an AD disease-control brain with antemortem clinical symptoms of frontotemporal dementia and an age-matched healthy control. We detect distinct laminar patterns of novel iron-laden glial pathology in both FTLD-Tau and FTLD-TDP brains. We find iron-positive ameboid and hypertrophic microglia and astrocytes largely in deeper GM and adjacent WM in FTLD-Tau. In contrast, FTLD-TDP presents prominent superficial cortical layer iron reactivity in astrocytic processes enveloping small blood vessels with limited involvement of adjacent WM, as well as more diffuse distribution of punctate iron-rich dystrophic microglial processes across all GM lamina. This integrated MRI/histopathology approach reveals *ex vivo* MRI features that are consistent with these pathological observations distinguishing FTLD-Tau and FTLD-TDP, including prominent irregular hypointense signal in deeper cortex in FTLD-Tau whereas FTLD-TDP showed upper cortical layer hypointense bands and diffuse cortical speckling. Moreover, differences in adjacent WM degeneration and iron-rich gliosis on histology between FTLD-Tau and FTLD-TDP were also readily apparent on MRI as hyperintense signal and irregular areas of hypointensity, respectively that were more prominent in FTLD-Tau compared to FTLD-TDP. These unique histopathological and radiographic features were distinct from HC and AD brains, suggesting that iron-sensitive T2*w MRI, adapted to *in vivo* application at sufficient resolution, may offer an opportunity to improve antemortem diagnosis of FTLD proteinopathies using tissue-validated methods.

## 1. INTRODUCTION

Frontotemporal lobar degeneration (FTLD) is an understudied family of age-associated neurodegenerative proteinopathies that encompasses a variety of progressive frontotemporal dementia (FTD) clinical syndromes and is the most common cause of young-onset dementia.^1^ Roughly 95% of FTLD is caused by one of two distinct proteinopathies: tauopathies (*i*.*e*., FTLD-Tau) or TDP-43 proteinopathies (*i*.*e*., FTLD-TDP).^2^ Despite the distinct postmortem neuropathology that distinguishes these proteinopathies microscopically, it is not currently possible to accurately detect and differentiate these pathologies during life.^3^ Indeed, while some frontotemporal dementia (FTD) clinical syndromes have group-level statistical associations with one of these proteinopathies, the majority of FTD syndromes do not reliably predict pathology.^3–9^ Moreover, there is currently no method to directly sensitize traditional MRI, or any other *in vivo* imaging technology, including molecular imaging, to the pathological features associated with underlying tau and TDP-43 inclusions in FTLD. In this report, we explore the novel combination of histopathology and T2*-weighted (T2*w) *ex vivo* 7 T MRI for the purpose of developing a more reliable, data-driven approach to diagnosis of pathology in FTLD spectrum disorders MRI-based studies of FTLD have focused on localizing neurodegeneration through quantification of cortical grey matter thinning and/or coherence of white matter fibers^10,11^. Both methods reveal group-level patterns of atrophy and disruption of neurocognitive networks that map to clinical symptoms.^12–15^ However, while gross regional patterns of neurodegeneration are associated at a group level with specific FTLD pathologies in autopsy data^6,8,9,16,17^, patient-level prediction of pathology using these methods remains elusive.^18^ Thus, *in vivo*, individual-patient level prediction of pathology is a major impediment and unmet need for development of disease-modifying therapies.^3,11^ In contrast, pathology-based studies of FTLD have suggested a variety of disparate microscopic features across tau-^19–21^ or TDP-43-mediated neurodegeneration^22,23^. However, microscopic studies are necessarily restricted in spatial scope due to the nature of histopathologic sampling and analysis.

*Ex vivo* MRI provides ultra-high-resolution imaging of fine structures over large sections of tissue. Previous work in other disorders has demonstrated significant correlations between T_2_*-weighted (T2*w) *ex vivo* MRI at 7 T and histopathology, particularly in the mapping of myelin and iron deposits in cortical laminae.^24–28^ Indeed, these two sources of contrast often overlap, as oligodendrocytes are a major source of non-heme iron in the brain due to the large metabolic demands for myelination.^29^ Joint MRI/pathology studies have particularly focused on Alzheimer’s disease (AD)^24,30–36^ and amyotrophic lateral sclerosis (ALS)^37–40^, both of which have been described producing specific and localized distributions of pathological intracortical iron. However, in FTLD, joint MRI/pathology has been largely limited to focused study of the basal ganglia.^41–43^

In this work we explore the combined use of histopathology and T2*w *ex vivo* 7 T MRI of whole brain hemispheres to 1) evaluate the sensitivity of T2*w MRI to detect and distinguish microscopic pathologic features of FTLD within cortical laminae and subjacent WM, and 2) demonstrate utility of MRI-guided histopathology to locate and typify focal pathologic features in FTLD. Previous histopathological studies suggest relative bilaminar distribution of tau inclusions^20,44,45^ and gliosis^46–49^ with greater WM degeneration^50,51^ in FTLD-Tau. In contrast, FTLD-TDP demonstrates greater relative upper cortical-layer TDP-43 inclusions^45,52,53^, degeneration and gliosis^23,54,55^. Therefore, we hypothesized T2w* MRI would reveal distinct laminar features of greater relative deep cortical layer and adjacent WM disease in FTLD-Tau compared to greater relative upper layer pathology in FTLD-TDP. Our findings’ laminar distribution were in concordance with this hypothesis, and, moreover, we describe below novel iron-rich patterns of gliosis in FTLD that highlight potential distinct mechanisms of neuroinflammation between proteinopathies.

## 2. MATERIALS AND METHODS

### 2.1 Patients and Neuropathological Diagnosis

Patients selected for study were evaluated at the Penn Frontotemporal Degeneration Center (FTDC). Clinical diagnosis was performed in a weekly, multidisciplinary consensus panel using published clinical research criteria for FTD.^4,5,56^ Autopsy was performed at the Penn Center for Neurodegenerative Disease Research (CNDR). Neuropathological diagnosis was performed using tissue from the non-scanned hemisphere by experienced neuropathologists (EBL, JQT) using well characterized antibodies^57^ and current diagnostic criteria.^58,2,59^

We examined 3 FTLD-Tau brains, and 2 FTLD-TDP brains, 1 Alzheimer’s disease (AD) brain and 1 age-matched healthy control brain to capture the heterogeneity of FTLD. Tauopathies can be subdivided into the prominent isoform of tau present in inclusions (i.e. 3-and 4-repeat, [3R, 4R]),^2^ and our sample contained both 3R and 4R pathologies. Similarly, FTLD-TDP can be subdivided into subtypes TDP-A, -B, and -C,^58^ and our sample contained both TDP-A and -C. Patient demographics and clinical characteristics are summarized in Table 1. All procedures were performed in accordance with Helsinki criteria and an informed consent procedure obtained in accordance with the University of Pennsylvania Institutional Review Board.

**Table 1.** Patient data and rating scores for pathological features of iron-rich gliosis in grey matter.

|  |  | Participant #1<br>HC | Patient #2<br>AD | Patient #3<br>TDP<br>C | Patient #4<br>TDP<br>A | Patient #5<br>Tau<br>GGT | Patient #6<br>Tau<br>PSP | Patient #7<br>Tau<br>PiD |
| --- | --- | --- | --- | --- | --- | --- | --- | --- |
| Age at Death (years) |  | 75 | 75 | 70 | 73 | 76 | 74 | 74 |
| Sex |  | F | M | M | F | M | M | M |
| Hemisphere sampled |  | R* | R | L | R | L | L | R* |
| Clinical diagnosis |  | HC | bvFTD | svPPA | bvFTD | naPPA/<br>PSPS | PSPS/<br>naPPA | bvFTD |
| Disease Dur. (years) |  | N/A | 6 | 7 | 14 | 4 | 7 | 6 |
| ABC |  | A0B1C0 | A2B3C3 | A1B0C0 | A0B2C0 | A2B0C0 | A2B0C3 | A1B0C0 |
| Brain Weight (g) |  | 1188 | 1258 | 1036 | 995 | 1316 | 1331 | 1217 |
| Post-Mortem Interval (hours) |  | 12 | 14 | 17 | 21 | 18 | 11 | 24 |
| Focal Region |  | MOT | ATP | ATP | IOFC | iMOT | MOT | DLPFC |
| Upper Layers I-III | Fe Glia | 0 | ++<br>Plaques/<br>Microglia | +++<br>Astrocyte<br>processes/<br>dystrophic<br>microglia | +++<br>Astrocyte<br>processes/<br>dystrophic<br>microglia | +<br>Ameboid<br>Microglia | +<br>Ameboid<br>Microglia | 0.5<br>Dystrophic<br>Microglia |
|  | 7T | 0 | Macro<br>Speckling | Band | Band | 0 | 0 | 0 |
| Deep Layers IV-VI | Fe Glia | 0 | +++<br>Plaques/<br>Microglia | +<br>Dystrophic<br>microglia | ++<br>Dystrophic<br>microglia | +++<br>Ameboid<br>Microglia | +++<br>Ameboid<br>Microglia | ++<br>Ameboid/<br>Dystrophic<br>Microglia |
|  | 7T | 0 | Macro<br>Speckling | Dots | Dots | Smudge | Smudge | Band |

Cells depict ordinal rating data (additionally color coded) for each pathology score: 0 (green) = no pathology, 0.5 (green) = rare pathology, + (yellow) = mild pathology, ++ (orange) = moderate pathology, +++ (red) = severe pathology or for 7T severity score
for MACRO SPECKLING = large hypointense speckling, BAND = hypointense banding pattern, DOTS= fine contrast inhomogeneity. Pt # = patient number, \* = frontal lobe imaged only, ABC = Amyloid Thal stage/ Braak AD Tau Stage/ CERAD plaque stage, bvFTD = behavioral variant of FTD, svPPA = semantic variant of primary progressive aphasia, naPPA = nonfluent agrammatic variant of primary progressive aphasia, GGT = globular glial tauopathy, PSPS = progressive supranuclear palsy syndrome, PiD = Pick's disease tauopathy; MOT = primary motor cortex; ATP = Anterior temporal pole; IOFC = inferior orbital frontal cortex; iMOT = inferior motor cortex; DLPFC = dorsolateral prefrontal cortex.

At the time of autopsy, one hemisphere was selected for standard neuropathological sampling from fresh tissue for diagnostics and frozen storage of remaining tissue as previously described,^57^ while the other hemisphere was immersed in 10% neutral buffered formalin for at least 30 days prior to imaging as below (Min=59, Max=626, Mean=187 days).

To harmonize with ongoing projects at our center, some hemisphere samples used for scanning were from the frontal lobe only and/or had 1.5 cm fresh samples taken prior to fixation for bilateral assessments of pathology in FTD, as previously described.^6^

### 2.2 *Ex vivo* 7 T MRI

Samples were placed in Fomblin (California Vacuum Technology; Freemont, CA), a proton-free fluid with volume magnetic susceptibility close to that of tissue. Samples were enclosed in either custom-build cylindrical holders or plastic bags, and then left to rest for at least two days to allow air bubbles to escape from the tissue. Depending on their size, samples were scanned using either a custom-built small solenoid coil or a custom-modified quadrature birdcage (Varian, Palo Alto, CA, USA) coil. These transmit/receive coils were attached to a two-channel transmit-receive adapter (Stark Contrast, Erlangen, Germany). The smallest coil was chosen that could hold each sample and then placed into our whole-body 7 T scanner (MAGNETOM Terra, Siemens Healthineers, Erlangen, Germany) using plastic shims under the coils to position the sample near isocenter.

MRI data were acquired with a 3D-encoded, 8-echo gradient-recalled echo (GRE) sequence with non-selective RF pulses. To maintain readout polarity and minimize distortions due to field inhomogeneity, each readout was followed by a flyback rephrasing gradient. The final echo was followed by an additional completely rephrased readout to measure frequency drifts. Each line of k-space was acquired with multiple averages sequentially before advancing to the next phase-encode step. Common parameters for the sequence were: 280 μm isotropic resolution, 25° flip angle, 60 ms repetition time (TR), minimum echo time (TE) 3.48 ms, echo spacing 6.62 ms, bandwidth 400 Hz/px. The field of view was adapted to each sample, and subsequently TRs and TEs were slightly modified based on the necessary readout duration. Total scan times were 8-10 hours for each sample.

Images were reconstructed using the vendor’s on-scanner reconstruction software which automatically corrected the global frequency drift, combined the signal averages in k-space, and produced magnitude images for each echo. After its MRI session, each sample was rinsed in formalin and then stored at room temperature in sealed bags for histopathological processing.

We determined that an echo time of roughly 20 ms provided strong T2*-weighted cortical laminar contrast while maintaining excellent SNR and minimizing local susceptibility induced drop-outs (*e*.*g*., due to air bubbles). We performed all subsequent analyses on these single-echo images. This choice of TE is consistent with some previous work using magnitude images directly,^25,60^ while slightly shorter than that chosen by others.^24,31,32,37^

At each pathology-sampled region (sampling process described below), MR images were rated by an experienced investigator (MDT). We quantified the relative contrast between pairs of neighboring structures using a range (−2, +2), where +2 indicates the first structure in the pair is substantially brighter than the second, -2 indicates the first structure in the pair is substantially darker than the second, and 0 indicates no apparent contrast between the structures.

### 2.3 Histopathologic Sampling

All patient samples for study were systematically sampled in the Penn Digital Neuropathology Lab using an atlas-based approach in key neocortical regions implicated in FTD,^6,7^ including orbitofrontal cortex (OFC, Brodmann area (BA) 11), anterior temporal cortex (ATC, BA 20 with the exception of patient #3, who only had tissue available for histology from adjacent BA 38 obtained fresh at autopsy), inferior prefrontal cortex (IPFC, BA 45) and primary motor (MOT, BA 4) known to be an area of high pathology in PSP.^21^ We additionally sampled primary somatosensory (SOM, BA 3) as a relative negative control region that is generally less-affected in FTLD.

Critically, in reviewing our MRI data, we noted unique focal cortical features outside of our standard sampling in some of our FTLD patients. To further investigate pathologic-imaging correlations, we additionally sampled these specific regions (see Supplementary Tables 1 and 4).

During both the atlas-and MRI-guided sampling, FreeView 6.0 (Athinoula A. Martinos Center for Biomedical Imaging, https://surfer.nmr.mgh.harvard.edu/fswiki/FreeviewGuide) was used to view the MR images and generate 3D surface models of the hemispheres on a bench-side computer. By matching gyral patterns on the hemisphere sample to the 3D surface model and MR images, the sampling team (DO, DJI, MDT) achieved consensus on the correspondence between imaging coordinates and histology sample locations. Samples, roughly 1.5 cm x 1.5 cm x 0.5 cm, were taken with cuts normal to the pial surface.

All samples were processed and embedded in paraffin, then sectioned into 6 um sections in the Penn Digital neuropathology lab for histological evaluation.^61^ Adjacent sections from each tissue sample were stained for healthy myelin using luxol-fast blue with hematoxylin and eosin to visualize GM (LFB),^50^ Meguro method for Perl’s Fe (iron) stain with DAB amplification,^62,63^ amyloid-beta (Nab228, CNDR),^64^ ferritin light chain (Abcam; Catalogue No ab69090), microglia (IBA-1; Santa Cruz Biotechnology Inc, Dallas, TX; Catalogue No sc-32725) and activated astrocytes (GFAP; Dako; Santa Clara, CA; Catalogue No Z0334). Moreover, sections for FTLD-Tau and AD were also immunostained for phosphorylated tau (AT8; Fisher, Waltham, MA; Catalogue No. ENMN1020)^65^ and FTLD-TDP for phosphorylated TDP-43 (p409.410; Protein Tech, Rosemont, IL; Catalogue No 66318-1-Ig)^66^ to identify protein inclusions characteristic of these disorders. All sections were counterstained with hematoxylin. To ensure specificity and reproducibility of chemical staining, sections were re-stained in three separate staining batches for LFB and iron stain.

Slides were reviewed by an experienced investigator (DJI) and rated for key histopathological features on a standardized 0-3 scale (i.e. none, mild, moderate, severe)^59^ including: 1) density of tau/TDP-43, glial and iron-rich pathology in supragranular GM upper (layers I-III), infragranular deep GM (infragranular IV-VI), juxtacortical WM enriched with U-fibers and adjacent relatively deep WM; 2) vacuolization and neuronal loss in upper and deep GM layers; and 3) the graded presence of intracortical WM (*i*.*e*., bands of Baillarger) and adjacent juxtacortical and relatively deeper WM in LFB and Fe-stained sections.

### 2.4 Data and Code Availability Statement

All MRI imaging data that support this study are openly available via Dryad at https://doi.org/10.5061/dryad.4tmpg4f8r. All histopathology data that supports this study are contained in the supplemental materials and tissue is available upon reasonable request from the authors, conditional on establishing a formal data sharing agreement with the University of Pennsylvania. No locally developed software was used in this study.

## 3. RESULTS

### 3.1 Summary of Observations

For brevity in the descriptions below, we will focus on the sample regions for each subject that had distinct MRI/histopathological findings compared to the control and/or other patient regions. These key regions of interest are summarized in Table 1. Please see Supplementary Tables 1-4 for full histopathological and imaging ratings.

### 3.2 Participant #1 — Healthy control

As expected, T2*w MRI showed dark WM with lighter cortical GM that, in turn, was generally darkest in lower layers, with one or two distinct tangential bands generally observed, depending on the cortical region (Supplementary Figure 1). Juxtacortical WM enriched with U-fibers generally showed a slight gradient of darker signal compared to deeper adjacent WM. Comparing with histology, these contrast differences were, as expected, produced by iron-containing myelinated fibers — radial fibers producing lower-layer cortical hypointensity and bands of Baillarger producing tangential hypointense “stripes”. There was also slightly greater density of iron-rich myelin in juxtacortical WM enriched for U-fibers around sulcal depths compared to relatively deeper WM.

**Figure 1.**
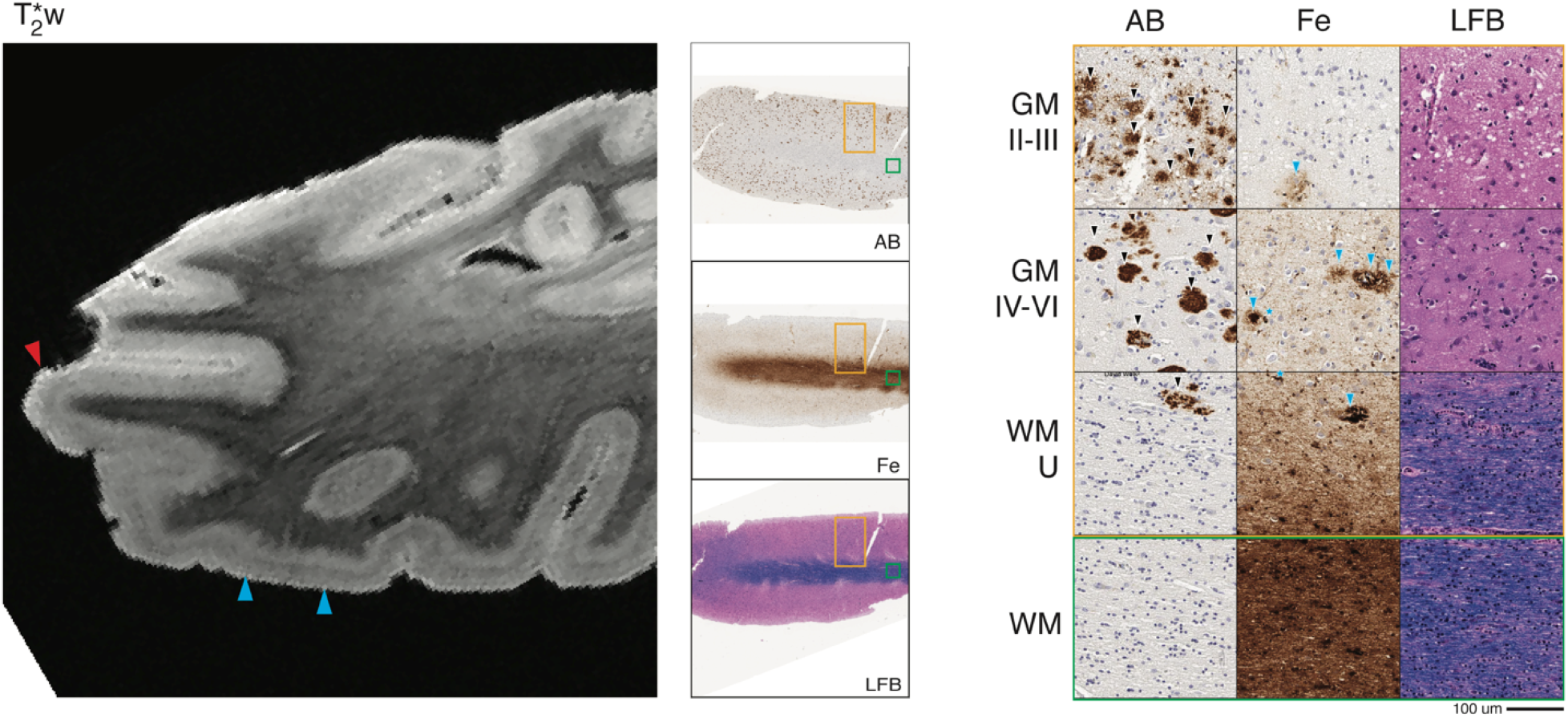
Alzheimer’s disease (patient #2) anterior temporal cortex MRI and pathology. *Left:* T2*w MRI of anterior temporal lobe (including sampled BA 20, red arrowhead shows sampled gyrus). *Center:* low-magnification (1x) view of tissue sample stained for amyloid-beta (AB), iron (Fe), and myelin (LFB). *Right:* high-magnification (20x) tissue sample stained for AB, Fe, and LFB (scale bar = 100 µm). Rows depict view in upper cortical layers (GM II-III), deep cortical layers (GM IV-VI), directly adjacent white matter enriched for cortical U-fibers (WM-U) and relatively deeper white matter (WM) from boxes outlined in low magnification view (orange box = cortical layers, green box = deep WM). There is widespread amyloid-beta plaque pathology (black arrowheads, AB) across cortical layers and in juxtacortical white matter along with severe neuronal loss across layers. T2*w MRI shows a widespread hypointense speckling pattern (blue arrowheads, T2*w) in mid-cortical layers which correlated iron deposits consistent in size and shape with neuritic senile plaques (blue arrowheads, Fe) and associated microglia (blue asterisks, Fe).

Consistent with the pathological staging data (Table 1) there was an absence of neurodegenerative disease-associated protein deposits in regions sampled. Examination of glial markers found mild IBA-1 reactivity throughout GM and WM largely in a “ramified” morphology of a quiescent resting type and only rare iron-or ferritin-reactive activated microglia with a rod-or ameboid-shaped morphology.^67,68^ GFAP staining revealed a “fibrous” morphology in sub-pial GM that seldom reached deeper than layer I GM and also more uniformly in WM, as described previously in healthy brain compared to a more activated “protoplasmic” morphology seen in neurodegeneration.^69,70^

### 3.3 Patient #2 – Alzheimer’s Disease, Behavioral-Variant Frontotemporal Dementia

Consistent with previous reports in amnestic AD, on MRI we noted a pattern of large hypointense speckling in GM in middle to lower cortical layers.^30–32,34,36^ This pattern was present throughout the neocortex and limbic cortex with relatively preserved patterns of GM lamination and WM architecture as found in the control sample. In our pathology-sampled regions, this speckling pattern was most prominent in the ATC (Figure 1).

Histopathologic examination of sampled tissue was consistent with neuropathological staging of high-level AD neuropathological change (Table 1) and revealed a high density of diffuse and neuritic amyloid-beta plaques and tau-positive tangles/threads largely confined to cortical GM.

Iron stain formed deposits that resembled a subset of iron-reactive amyloid-beta plaques and associated microglia in activated morphology (Figure 1) that were most pronounced in ATC and OFC. The distribution and intensity of myelin and iron stain also indicated relative preservation of intracortical myelin and adjacent WM throughout most regions sampled, although there was reduced density in ATC.

### 3.4 Patient #3 – FTLD-TDP Type C, Semantic Variant PPA

On MRI, atypical signal was largely localized in the ATC, consistent with the patient’s clinical syndrome of svPPA.^4^ ATC showed substantial atrophy and a hypointense “band” in the upper layers of the cortex, along with mild diffuse dot-like hypointense speckling across the entire cortical depth with loss of distinct cortical GM lamination seen in the control sample. A thin band of juxtacortical WM also showed overall hypointense signal compared to relative hyperintense signal in the focal area of relative deep WM to this cortical region. This regionally hyperintense signal in WM was accompanied with diffuse hypointense striations on MRI (Figure 2).

**Figure 2.**
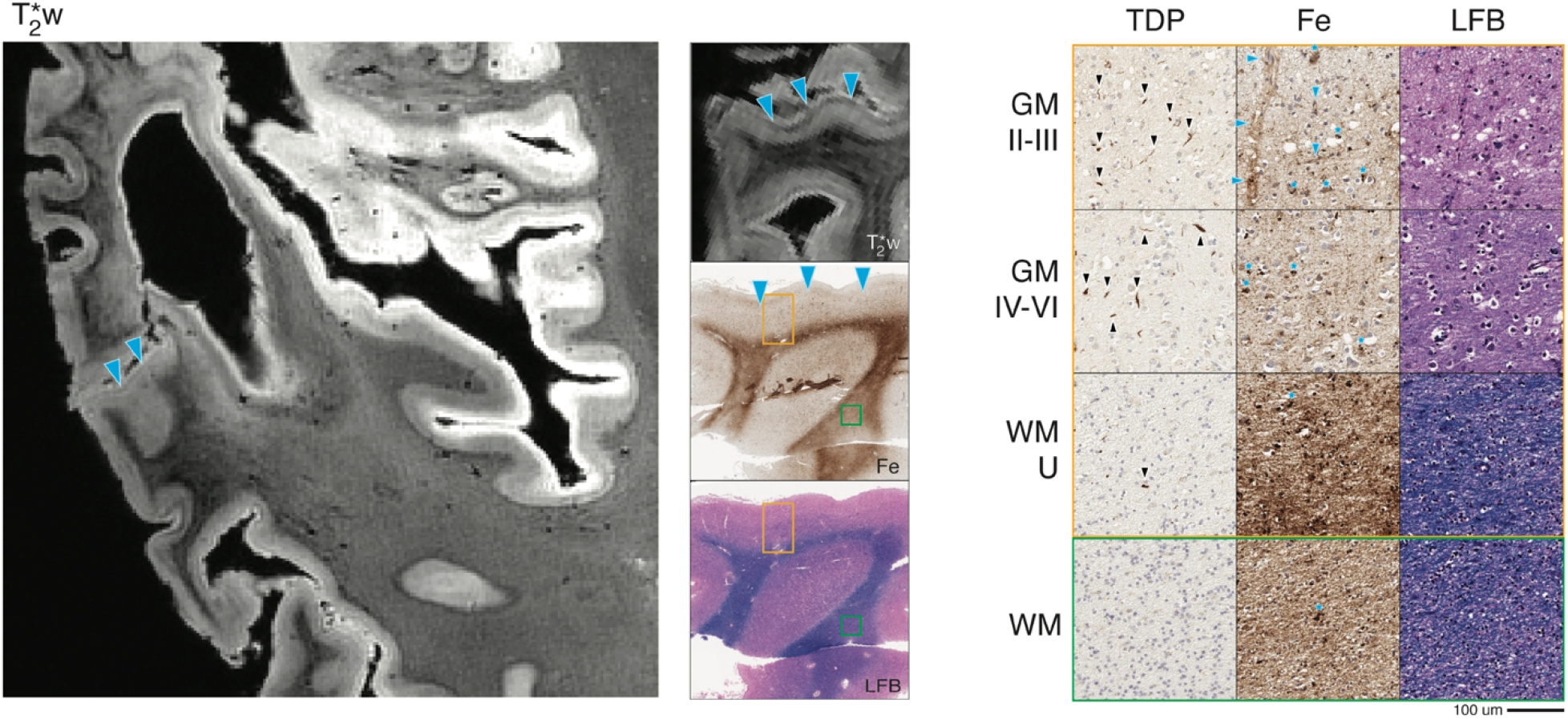
FTLD-TDP type C (patient #3) anterior temporal cortex MRI and pathology. *Left:* T2*w MRI of anterior temporal lobe (including sampled BA 38). *Center:* low-magnification (1x) view of tissue sample stained for iron (Fe) and myelin (LFB), with corresponding T2*w MRI slice. *Right:* high-magnification (20x) tissue sample stained for TDP-43 pathology (TDP), Fe, and LFB (scale bar = 100 µm). Rows depict view in upper cortical layers (GM II-III), deep cortical layers (GM IV-VI), directly adjacent white matter enriched for cortical U-fibers (WM-U) and relatively deeper white matter (WM) from boxes outlined in low magnification view (orange box = cortical layers, green box = deep WM). There is prominent TDP-43 pathology (black arrow heads, TDP) across cortical layers and most abundant in upper layers along with severe neuronal loss and vacuolization (LFB). T2*w MRI shows a hypointense upper-layer band (blue arrowheads, T2*w) and diffuse speckling which correlated with iron deposits resembling astrocyte processes surrounding capillaries (blue arrowheads, Fe) and activated microglia (blue stars, Fe). Outside of the upper-layer band, there is an absence of intracortical contrast on MRI (T2*w), and rare myelin fibers on histology (LFB). Adjacent white matter signal is heterogenous on MRI (T2*w) corresponding to mild reduction in relative deep white matter myelin (LFB) and occasional white matter iron rich glia (blue arrowheads, Fe).

Histopathological analysis of ATC showed a high density of TDP-43 positive long-dystrophic neurites that was most prominent in upper layers and associated with relatively greater superficial cortical layer neuronal loss and vacuolization. TDP-43 pathology was largely absent in WM, similar to previous reports for FTLD-TDP type C.^22^ The remainder of regions had mild or absent TDP-43 and neurodegeneration.

Corresponding to the observed upper layer hypointense band and diffuse cortical speckling seen on MRI, iron staining in ATC revealed a high density of staining in cortical layer II, consisting of a mixture of dot-like stippling pattern of bead-like dystrophic appearing processes and additional prominent reactivity of cellular processes surrounding small blood vessels (Figures 2, 7). Examination of adjacent sections for glial markers suggested that these iron-rich structures correspond to diffuse punctate and tortuous IBA-1-positive dystrophic microglial processes and GFAP-positive astrocytic processes enveloping small vessels, respectively. Moreover, there were moderate amounts of iron-rich cellular structures resembling rod-shaped or ameboid microglial morphologies in layers III-VI and occasionally in juxtacortical WM that corresponded to activated microglia visualized by IBA-1 immunostaining (Figure 7). Histologically, WM staining in ATC showed scant intracortical myelin and mild relative loss of adjacent WM compared to control tissue.

### 3.5 Patient #4 – FTLD-TDP Type A, Behavioral Variant FTD

On MRI, both ATC and OFC showed atypical diffuse dot-like speckling throughout the cortex, including upper layers. There was focal hyperintense signal in adjacent WM in ATC with a gradient of darker signal in juxtacortical WM and some scant WM striations of hypointense signal. Similar to the TDP C case (patient #3), we found an upper-cortical hypointense band on MRI, here in a lateral region of OFC (BA 47) (Figure 3).

**Figure 3.**
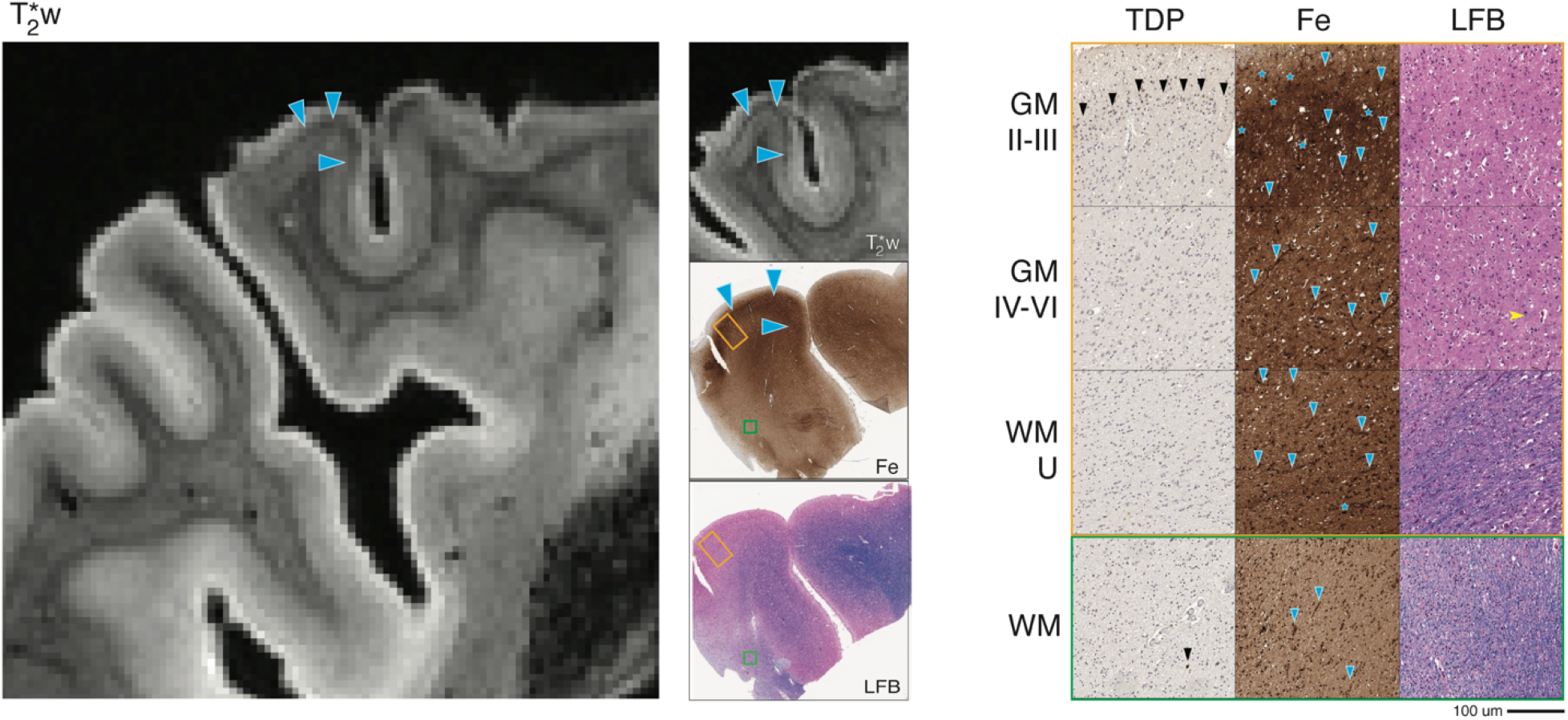
FTLD-TDP type A (patient #4) lateral orbitofrontal MRI and pathology. *Left:* T2*w MRI of lateral orbitofrontal region (including sampled BA 47). *Center:* low-magnification (1x) view of tissue sample stained for iron (Fe) and myelin (LFB), with corresponding T2*w MRI slice. *Right:* high-magnification (20x) tissue sample stained for TDP-43 pathology (TDP), Fe, and LFB (scale bar = 100 µm). Rows depict view in upper cortical layers (GM II-III), deep cortical layers (GM IV-VI), directly adjacent white matter enriched for cortical U-fibers (WM-U) and relatively deeper white matter (WM) from boxes outlined in low magnification view (orange box = cortical layers, green box = deep WM). There is prominent TDP-43 pathology largely restricted to upper cortical layers (black arrow heads, TDP) along with severe neuronal loss and vacuolization (LFB). T2*w MRI shows a hypointense upper layer band and diffuse speckling (blue arrow heads, T2*w) which correlated with iron deposits resembling astrocyte processes surrounding capillaries (blue arrow heads, Fe) and iron-rich glia (blue stars, Fe). There were sparse deep fibers in the deeper cortical layers (yellow notched arrow head, LFB). Adjacent white matter signal was heterogenous on MRI, corresponding to mild reduction in relative deep white matter myelin (LFB) and occasional white matter deposits and gliosis (blue arrowheads, Fe).

In ATC and OFC, histopathology showed highest TDP-43 pathology in cytoplasmic inclusions and diffuse neurites, accompanied by severe upper layer neuronal loss and vacuolization in regions sampled, consistent with known pathological and regional patterns of FTLD-TDP A.^22,58^ There were also mild TDP-43 inclusions in WM oligodendrocytes in these regions, while the remainder of regions sampled had scant or absent TDP-43 and minimal or no neurodegeneration of GM or WM.

The MRI-guided lateral OFC sample showed a similar pattern on histopathology, with more pronounced iron-rich astrocytic processes enveloping small vessels in layers II-III and overall darker appearance of the neuropil, particularly in upper layers (Figure 3), similar to patient #3 with FTLD-TDP C. Iron-reactive pathology corresponded to a high level of GFAP-reactive astrocytes highlighting small capillaries and beaded dystrophic IBA-1 reactive microglial processes (Figure 7). Intracortical myelin was mildly reduced, as was adjacent WM. In the standard sampled OFC and ATC there was a similar pattern with less prominent upper-layer iron-rich gliosis and ATC had more prominent iron-positive hypertrophic ameboid microglia.

### 3.6 Patient #5 – 4R-Predominant Globular Glial Tauopathy, Non-Fluent/Agrammatic PPA

MRI showed prominent abnormal signal in MOT, which showed frank irregular hypointense “smudges” within the deep layers of cortical laminae. Adjacent WM showed large irregular hypointense striations with vessel-like shapes. These findings were most pronounced in the inferior aspect of the motor cortex near the areas involved with motor speech, matching the patient’s clinical presentation of naPPA.^4^ We performed MRI-guided sampling of an additional inferior area of motor cortex with most prominent imaging features (iMOT) (Figure 4).

**Figure 4.**
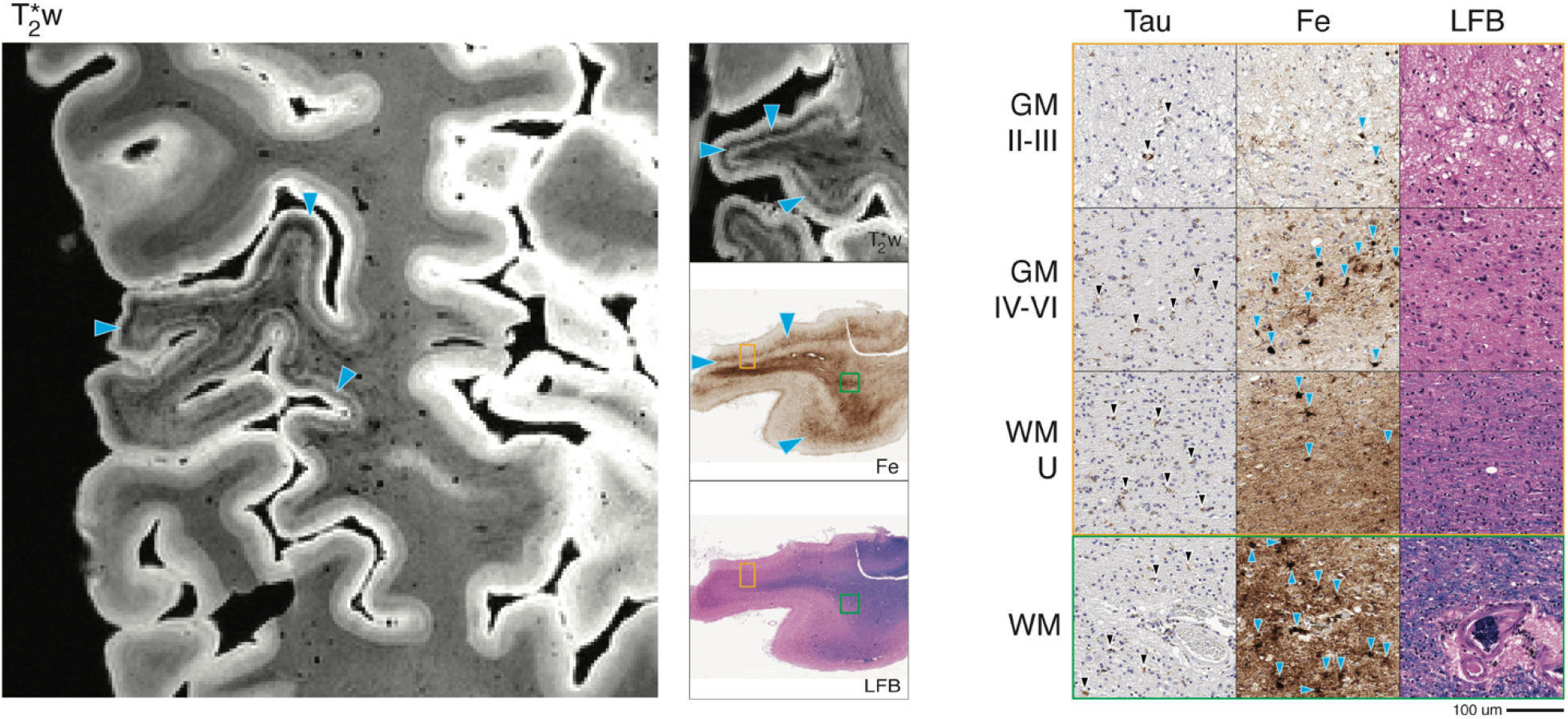
FTLD-Tau GGT 4R tauopathy (patient #5) primary motor MRI and pathology. *Left:* T2*w MRI of the inferior aspect of the primary motor region (including sampled BA 4). *Center:* low-magnification (1x) view of tissue sample stained for iron (Fe) and myelin (LFB), with corresponding T2*w MRI slice. *Right:* high-magnification (20x) tissue sample stained for tau pathology (Tau), Fe, and LFB (scale bar = 100 µm). Rows depict view in upper cortical layers (GM II-III), deep cortical layers (GM IV-VI), directly adjacent white matter enriched for cortical U-fibers (WM-U) and relatively deeper white matter (WM) from boxes outlined in low magnification view (orange box = cortical layers, green box = deep WM). There is widespread tau pathology (black arrowheads, Tau) across cortical layers and in white matter along with severe neuronal loss and vacuolization across layers (LFB). T2*w MRI shows a large irregular hypointense band (blue arrow heads, T2*w) across mid to lower cortical layers. This coincided on histopathology with clusters iron deposits resembling hypertrophic appearing microglia (blue arrow heads, Fe). There was no apparent MRI signal corresponding to the band of Baillarger and adjacent white matter was striated with large hypointensities near blood vessels. This pattern corresponded on pathology to an absence of intracortical white matter and severe myelin loss in adjacent white matter, along with large clusters of iron deposits resembling glia surrounding larger blood vessels (blue arrow heads, Fe).

This patient displayed 4R-predominant pathology in tau-positive “globular” oligodendrocytic tau inclusions in GM and WM that was most prominent in the corticospinal tract as well as astrocytic tau pathology, tangles and threads in all layers of cortical GM, consistent with globular glial tauopathy type II.^71^ Tau pathology and resultant neurodegeneration were severe and most prominent in primary MOT and iMOT regions, while milder in IPFC and OFC and negligible in SENS. There was also mild to moderate amount of co-morbid diffuse amyloid beta plaque (stage A2, C0-Table 1) most concentrated in OFC and IPFC with rare iron-reactivity resembling plaques for these regions only.

Corresponding to irregular hypointense signal on MRI, iron stain of MOT and iMOT revealed a central band (approximately layers III-V) of dense, highly-reactive iron-positive clusters resembling ameboid and hypertrophic microglia (Figure 4). There were also lighter iron-reactive deposits corresponding to astrocyte morphologies in mid-cortical layers and in sub-pial areas of layer I. Adjacent tissue stained for glial markers confirmed the presence of large ameboid-type microglia in mid to deeper layers of GM and throughout WM, as well as a high density of weakly GFAP-positive reactive protoplasmic astrocytes with some dystrophic features throughout GM layers with a relative depletion of fibrous astrocytes in WM compared to the control sample (Figure 7). Myelin stain showed minimal existing intracortical myelin and moderate loss of WM in juxtacortical and relative deep areas of adjacent WM. Rare diffuse amyloid plaques that did not correspond to iron stain were noted in MOT and iMOT, and were spatially distinct from the area of novel iron-rich glial pathology in this sample (Supplementary Figure 2). Iron staining of subjacent WM showed similar patches of high density of iron-positive microglia, often in focal clusters surrounding large blood vessels. Vessel walls appeared thickened with enlarged surrounding space and mild extracellular hemosiderin deposits but no evidence of amyloid angiopathy in these large vessels in relatively deeper WM (Supplementary Figure 2).

### 3.7 Patient #6 – 4R-Predominant PSP Tauopathy, Steele-Richardson Progressive Supranuclear Palsy Syndrome

MRI results in our standardized sampling regions identified a distinct pathological feature in MOT, which was notable for a large irregular hypointense “smudge” in the middle layers of cortex with some irregular extension of mild hypointensity into the subjacent WM relative to nearby gyri (Figure 5), similar to GGT. This irregular hypointense band partially obscured the signal from intracortical WM. Finally, there was large-sized speckling across the cortical layers in ATC and IPFC which were reminiscent of that seen in the AD patient.

**Figure 5.**
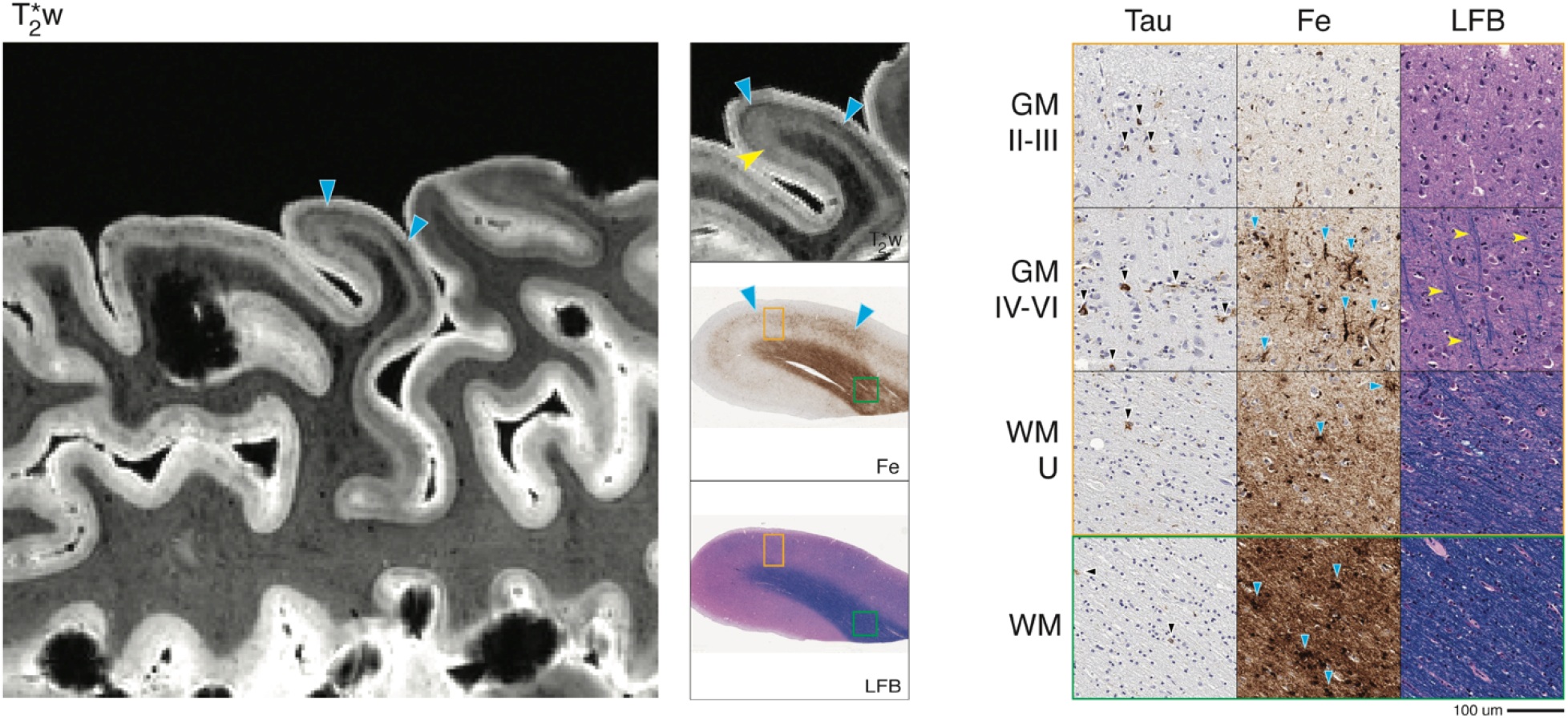
FTLD-Tau PSP 4R tauopathy (patient #6) primary motor MRI and pathology. *Left:* T2*w MRI of the primary motor region (including sampled BA 4). *Center:* low-magnification (1x) view of tissue sample stained for iron (Fe) and myelin (LFB), with corresponding T2*w MRI slice. *Right:* high-magnification (20x) tissue sample stained for tau pathology (Tau), Fe, and LFB (scale bar = 100 µm). Rows depict view in upper cortical layers (GM II-III), deep cortical layers (GM IV-VI), directly adjacent white matter enriched for cortical U-fibers (WM-U) and relatively deeper white matter (WM) from boxes outlined in low magnification view (orange box = cortical layers, green box = deep WM). There is widespread tau pathology (black arrowheads, Tau) across cortical layers and in white matter along with moderate neuronal loss across layers (LFB). MRI showed a large irregular hypointense band across mid-to-lower cortical layers (blue arrowheads, T2*w); on histopathology this correlated to clusters of iron deposits resembling hypertrophic appearing microglia (blue arrowheads, Fe). There was some preserved shading on MRI (yellow notched arrowhead, T2*w) that was partially obscured by the pathological iron signal; this pattern corresponded to the preservation of deep cortical layer myelin (yellow notched arrowheads, LFB). Adjacent white matter showed patchy hypointensities, which corresponded on pathology to the mild reduction in myelin (LFB) and additional large clusters of iron deposits resembling glia (blue arrowheads, Fe).

Histopathology was typical for PSP, with tau-positive tangles and tufted astrocytes throughout cortical layers and tau-positive threads and coiled bodies in adjacent WM that was most severe in MOT.^21^ This patient had moderate AD plaque co-pathology (Table 1) and a subgroup of amyloid-plaques were iron-reactive in ATC and IPFC, correlating with our observation on MRI of “large-size speckling” in the ATC and IPFC. In contrast, amyloid-beta plaque co-pathology was mild in MOT, and spatially distinct from the band of unique iron-positive hypertrophic microglia in this sample (Supplementary Fig 2) described below.

Corresponding to the deep-layer hypointensity on MRI, iron stain showed clustered deposits resembling activated microglia in layers III-V and adjacent WM, similar to GGT (Figure 5). These clusters were occasionally associated with large blood vessels in GM and WM. There were less distinct astrocyte morphologies seen on iron stain, and adjacent tissue stained for GFAP showed mild reactivity in mid-layers and similar findings to controls in WM (Figure 7). Myelin histology revealed preserved deep layer cortical myelin and adjacent WM..

### 3.8 Patient #7 – 3R-Predominant Tauopathy, Pick’s Disease, Behavioral-Variant Frontotemporal Dementia

MRI identified OFC and IPFC as abnormal among our standard sampling. In both of these regions, and diffusely thought the frontal cortices, there was overall WM hyperintensity relative to posterior healthy appearing WM associated with MOT and SENS, with the effect most pronounced in OFC. In frontal association regions there was an atypical lack of contrast from deep and upper cortical layers as seen in the control brain. We located an additional focal area in the dorsolateral midfrontal cortex (DLPFC, BA9) with mid-to-deep cortical hypointense band which we sampled for additional histopathologic analysis (Figure 6).

**Figure 6.**
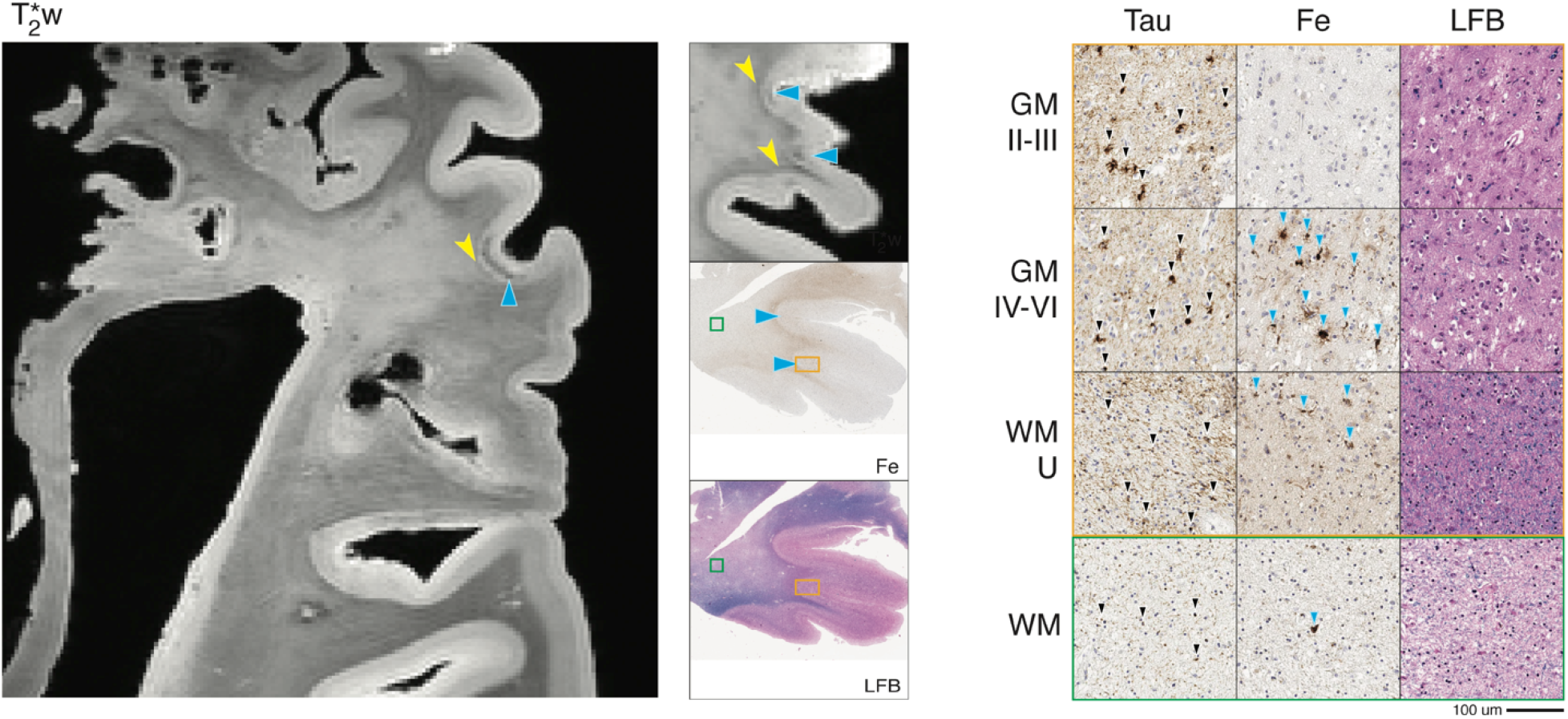
FTLD-Tau Pick’s disease 3R tauopathy (patient #7) dorsolateral prefrontal MRI and pathology. *Left:* T2*w MRI of a dorsolateral prefrontal region (including sampled BA 9). *Center:* low-magnification (1x) view of tissue sample stained for iron (Fe) and myelin (LFB), with corresponding T2*w MRI slice. *Right:* high-magnification (20x) tissue sample stained for tau pathology (Tau), Fe, and LFB (scale bar = 100 µm). Rows depict view in upper cortical layers (GM II-III), deep cortical layers (GM IV-VI), directly adjacent white matter enriched for cortical U-fibers (WM-U) and relatively deeper white matter (WM) from boxes outlined in low magnification view (orange box = cortical layers, green box = deep WM). There is widespread tau pathology (black arrow heads, Tau) across cortical layers and in white matter along with severe neuronal loss across layers (LFB). MRI shows a large irregular hypointense band (blue arrowheads, T2*) in mid-to-lower cortical layers; this coincided on pathology with clusters of iron-rich hypertrophic appearing microglia and astrocytic profiles (blue arrowheads, Fe). There were also clusters of iron rich glia in juxtacortical WM (blue arrowheads, Fe) associated on MRI with relative hypointensity of juxtacortical WM (yellow notched arrowheads, T2*w) compared to the hyperintense deeper WM.

**Figure 7.**
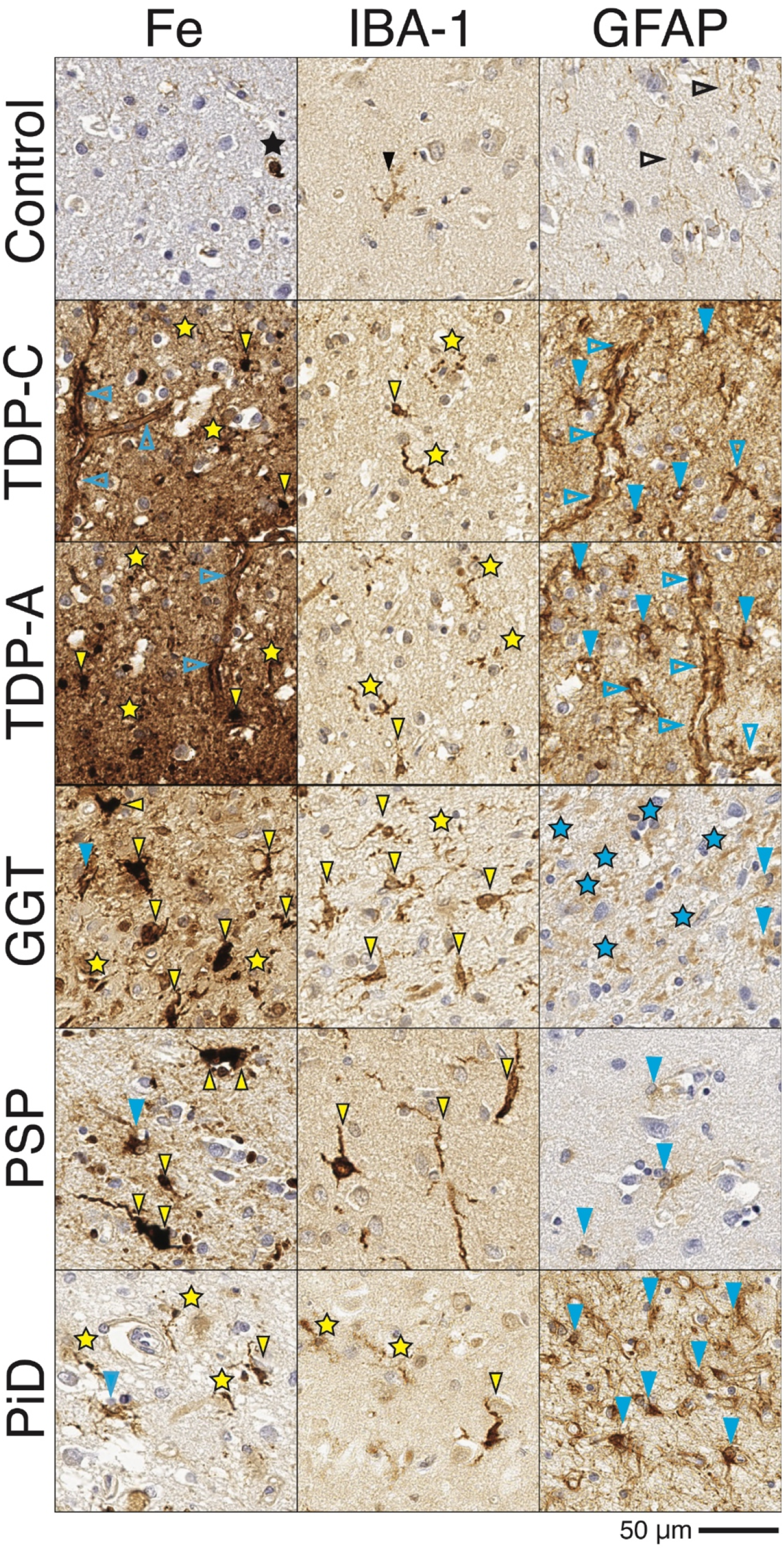
Patterns of iron-rich glia in FTLD-TDP and FTLD-Tau groups. Photomicrographs depict representative images from iron-stained sections with abnormal T2* signal ex vivo in FTLD samples and semi-adjacent sections immunostained for microglia (IBA-1) and activated astrocytes (GFAP). The healthy control (HC) had iron-reactivity largely restricted to oligodendrocytes (black star). Glial immunostaining in HC shows mild amounts of microglia in ramified non-reactive morphology (solid black arrowhead) and mild diffuse GFAP+ fibrous astrocytic processes in upper cortical layers (open black arrow heads). The FTLD-TDP patients (TDPA and TDPC) had shared features of relative upper layer (layers II-III) iron in processes surrounding capillaries (open blue arrowheads) as well as diffuse speckling of the neuropil (yellow stars) corresponding to a high level of GFAP reactive astroglial processes (open blue arrowheads) and dystrophic processes (yellow stars) of IBA-1 reactive microglia (solid yellow arrowheads), respectively. In contrast, FTLD-Tau patients had prominent clusters of hypertrophic “ameboid” and “rod-like” appearing iron-rich profiles (solid yellow arrowheads) that were most conspicuous in middle to deeper cortical layers and corresponded to IBA-1 reactive microglia in hypertrophic morphologies (solid yellow arrowheads). The GGT sample also showed dystrophic processes (yellow stars) on iron-stain and IBA-1, with prominent fragmented dystrophic GFAP+ processes (blue stars) throughout. In contrast, PSP showed moderate level of astrogliosis (open blue arrowhead) that were less similar in morphology to adjacent iron-stained cellular structures. The Pick’s disease (PID) 3R tauopathy had similar, but relatively isolated, clusters of hypertrophic iron-rich microglia (solid yellow arrowheads) in deep grey matter and adjacent juxtacortical white matter that corresponded to activated IBA-1 morphologies and degenerating processes (yellow stars). There were also occasional iron-rich morphologies similar to GFAP-reactive activated astrocytes (solid blue arrowheads) which were more prominent in PID than in the 4R tauopathies, but in contrast to FTLD-TDP, were seldom iron-rich. Scale bar = 50 µm.

Histopathologic examination identified tau-positive Pick bodies in neurons and tau-positive ramified astrocytes throughout the cortical layers and associated with severe neuronal loss typical for Pick’s disease in the OFC, IPFC, and DLPFC.^20^ This was accompanied by severe tau pathology in the form of WM threads and oligodendrocytes in these regions. In contrast, tau pathology and neurodegeneration were minimal in primary MOT and SENS cortex.

Iron staining in DLPFC found prominent clusters of iron-positive activated microglia and dystrophic processes in mid to deep GM layers (layers III-VI) and juxtacortical WM and less prominent iron-positive astrocytes in deep GM. There were also scattered reactive iron-positive microglia in deeper adjacent WM. Interestingly, glial staining in adjacent tissue revealed a relative depletion of microglia in DLPFC with clusters of microglia with ameboid profiles in a similar spatial distribution to iron stain and severe widespread GFAP reactivity in astrocytes and their processes in GM and WM (Figure 7). In contrast, OFC and IFC, which had similar tau pathology and neurodegeneration to DLPFC, had only scant iron deposits resembling glia. Thus, the observed hyperintensity of cortex on MRI appeared to derive from both the lack of cortical myelin and rarity of iron-rich glia. Finally, iron staining in OFC, IPFC and DLFPC also reflected the absence of cortical myelin and reduced adjacent WM integrity correlating with MRI findings of indistinguishable cortical lamination and hyper intense signal in adjacent WM in these regions (Figure 6).

## 4. DISCUSSION

We used an integrated, *ex vivo* MRI and histopathology approach to study diverse FTLD clinical and pathological subtypes, and contrasted these with AD and healthy brain samples. This enabled novel findings of distinct upper layer hypointense bands and diffuse speckling on T2*w MRI which corresponded to both iron-rich astrocytic processes surrounding small blood vessels and dystrophic (as opposed to hypertrophic) patterns of microglia, respectively in FTLD-TDP. In contrast, these patterns were not observed in FTLD-Tau and instead we found large irregular hypointense signal in deep cortical layers and adjacent WM on T2*w MRI that correlated with iron-rich hypertrophic microglia. Moreover, there was loss of normal cortical myelination patterns and hyperintense signal in adjacent WM corresponding to severe WM degeneration particularly evident in the 3R tauopathy PID brain. Thus, our findings suggest potential divergent mechanisms of neuroinflammation and resultant neurodegeneration between specific pathological forms of FTLD that may be detectible during life. We also demonstrate the utility of *ex vivo* MRI to guide histopathologic work needed to study these novel patterns of gliosis outside of traditional histopathological sampling optimized for AD^59^

### 4.1 Focal iron-positive gliosis within the cortical laminae and adjacent white matter in FTLD proteinopathies

In both FTLD-TDP cases we found an irregular upper-layer hypointense band within the cortex on MRI, corresponding to iron deposits and GFAP-positive astrocytic processes enveloping small blood vessels. Additionally, hypointense speckling in the cortex of diseased regions was observed on MRI throughout cortical layers that correlated on histopathology with dystrophic microglial processes labelled by IBA-1 (Figures 2, 3, 7).

These findings are similar to previous reports of iron-rich gliosis in motor cortex of ALS with TDP-43 proteinopathy, but contrast with the laminar distribution described in ALS as mid-to-deep layers.^37,72^ Our study did not include ALS or TDP-B, both of which have greater relative deep layer pathology compared to TDP-A and -C,^58^ which may contribute to this discrepancy. Nonetheless, these patterns were distinct from those seen in FTLD-Tau patients in our cohort.

On MRI, the 4R tauopathy cases (*i*.*e*., GGT and PSP, patients 5, 6) had prominent mid-to-deep layer irregular hypointense “smudges” in GM, and vessel-shaped hypointensities in nearby WM. On pathology, these corresponded to clusters of iron-rich activated microglia often surrounding large vessels (Figures 4, 5, 7). While the gross anatomical location of a high burden of pathology in primary MOT in these tauopathies^21,71^ partially overlaps with regional patterns of pathology in ALS,^37,73^ the radiographic and pathological features of these FTLD-Tau cases are distinct in the addition of prominent iron-rich gliosis in adjacent WM. Indeed, a previous report of GGT histology also finds similar pattern of iron-rich gliosis in deep GM and adjacent WM.^48^

In PiD, the 3R tauopathy (patient 7), there were shared features with 4R tauopathies of focal, deep-layer hypointensity on MRI, occurring in this case in DLPFC. On pathology, this was associated with a mixture of activated astrocytes and dystrophic microglia processes that were also evident in juxtacortical WM (Figures 5, 7).

Despite similar clinical symptomatology, our FTLD samples had distinct MRI and pathological features from our AD patient. Several studies in AD using *ex vivo* MRI and pathology have described non-homogenous appearance of cortex with hypointense speckling in layers III-V that correlates with varying amounts of iron-rich plaque and associated microglia on histology.^30,31,36^ Similar to the previous studies, we observed this mid-layer speckling pattern in our AD subject with antemortem bvFTD diagnosis, which was associated on histology with a subset of neuritic plaques and activated microglia forming morphologies and distributions concordant with iron deposits in mid cortical layers (Figure 2). Interestingly, the ATC, a region implicated in bvFTD,^74^ was an area of high iron deposits and corresponding amyloid-beta plaque pathology in this patient.

### 4.2 Histological analysis of iron-positive glia suggests disparate mechanisms of iron homeostasis and inflammation in AD and FTLD subtypes

Our histological analysis found only a subset of activated glia appear to sequester iron (Figure 7). Alterations of iron homeostasis have been linked to neurodegeneration as deficits in autophagy may lead to lysosome-mediated release of ferric iron from ferritin in degenerating cells (reviewed in detail elsewhere^75–77^). Glial activation may also contribute to neurodegeneration in AD, FTLD, and related disorders via non-cell autonomous mechanisms altering the microenvironment of the brain from chronic neuroinflammation.^78,79^

The large iron-rich ameboid morphologies seen in areas of high-pathology in 4R tauopathies suggest the pro-inflammatory state may involve iron sequestration from phagocytosis of cellular debris.^80^ In particular, the iron-stained features consistent with ameboid microglia in WM were uniquely localized near larger blood vessels in our 4R tauopathy samples. Indeed, ameboid activated microglia have been described in 4R tauopathies^48,81^ and similar findings of a large burden of perivascular phagocytic microglia were reported in a hereditary tauopathy.^82^ While we saw similar iron-rich ameboid morphology in TDP samples, these were less common and more focally only in the ATC of FTLD-TDP subtype A patient #4.

In contrast to the ameboid form, we also noted cases where iron deposits had a beaded appearance that morphologically resemble senescent microglia.^83,84^ This was particularly evident in FTLD-TDP (patients #3 and #4), and focally in deep layers of the DLPFC of the PiD sample (patient #7). These findings align with previous reports of prominent upper layer microglial activity in forms of FTLD-TDP^54,55^ while PiD shows a variable but more uniform distribution across layers.^49,54^

More generally, in our FTLD-TDP samples we found localized iron deposits and corresponding microglia in focal areas of cortical degeneration, in contrast to the large-vessel patterns seen in large areas of WM in 4R tauopathies. Indeed, activatedmicroglia have been associated with focal areas of cortical degeneration in FTLD-TDP suggesting an association of WM microglia with Wallerian degeneration of axons from neuronal loss.^85^

GFAP-reactive astrocytes are normally found in healthy white matter in a fibrous morphology,^69^ as seen in our control sample. However in our GGT and PiD sample (patients #5 and #7), there was a conspicuous absence of these fibrous astrocytes in WM and instead there was fragmentation of GFAP-positive processes in GGT (Figure 7), similar to previous reports of astrocytic degeneration in FTLD-Tau.^86,87^ FTLD-Tau has prominent tau pathology within astrocytes and this mechanism could contribute to the patterns observed here.

In contrast, FTLD-TDP inclusions are seldom localized within astrocytes^88^ and we did not observe degeneration of astrocytes in these samples. Instead, in FTLD-TDP we found prominent GFAP-reactive astrocytic morphologies in diseased regions, highlighted by iron deposits resembling astrocytic processes surrounding small vessels in upper cortical layers (Figures 2, 3, 7). Similar to previous reports,^46,47,49^ we also observed high levels of GFAP-positive gliosis in PiD across all cortical layers but, in contrast to FTLD-TDP, these cellular structures were largely not visualized by iron staining, suggesting a potential unique process of astrocyte-mediated iron dysregulation in TDP-43 proteinopathies. This is consistent with the likely contribution of astrocytes to iron homeostasis via active transport uptake of peripheral iron.^89^

### 4.3 Differential depletion of healthy myelin across FTLD, AD, and control

Healthy myelin, as observed in our control tissue, has distinct laminar distribution across cortical regions (i.e. myeloarchitecture) with most prominent intracortical myelin found in deeper layers (Nieuwenhuys et al., 2015). Moreover, previous work has found various myelination patterns in other primary and secondary associative cortices (Fracasso et al., 2016) that we largely recapitulate here (Supplementary Figure 1). In our FTLD samples, we found novel evidence of striking deviation from these established patterns of myelination in healthy cortex and adjacent WM.

The most obvious loss of intracortical myelin was seen in our PiD sample, appearing on MRI as widespread homogenous hyperintense signal in frontal association cortex, corresponding histologically to obliteration of deep-layer myelin and severe neuron loss throughout all layers (Figure 6). A similar but more focal pattern of intracortical myelin loss was seen in the GGT patient in primary motor cortex (Figure 4) while the band of Baillarger in the PSP sample (Figure 5) was relatively preserved. Heterogeneity observed among tauopathies could be attributed to tau isoform type (3R vs 4R), individual differences in rate of progression, disease duration, and/or the relative minimal distribution of PSP tauopathy in cortical regions compared to brainstem and subcortical regions not examined here.^21^ In FTLD-TDP samples, there was also evidence for degeneration of intracortical WM that correlated with severe neuronal loss and was most prominent in upper layers II-III (Figures 2, 3). Thus, neuronal loss and WM pathology may contribute to loss of myelination in our samples.

This differed from the pattern of cortical lamination in our atypical AD patient. While largely preserved on histology, cortical lamination was obscured in many locations on MRI by the non-homogeneous hypointense speckling associated with iron deposits resembling and spatially correlated with a subset of neuritic plaques and associated glia (Figure 2). One previous study found a subset of advanced AD patients, largely with young age of onset, to have an abnormal mid-layer band pattern on T2*w MRI corresponding to increased size and disorganization of intracortical myelin.^32,90^ We did not observe this pattern in our patient sample, which, had a relative older age at onset (75) which could account for this discrepancy.

### 4.4 Limitations and future work

The strong correlation between *ex vivo* T2*w MRI contrast and the presence of iron in our pathology analyses suggests that iron-based contrast in reactive/degenerating glia and healthy myelin is the largest, most consistent effect across our samples. Other sources of T2*w have been suggested. For example, diamagnetism of tau and beta amyloid inclusions has been quantified *in vitro* and in transgenic mice, finding that tau protein inclusions were significantly diagmagnetic at echo times in the range used in our study, with beta amyloid showing a smaller effect.^33^ In our sample this does not appear to produce significant T2*w contrast separate from iron, as exemplified in our PiD sample (patient #7), where OFC and IPFC had substantial tau inclusions but minimal iron, and was generally homogenous and hyperintense on MRI (Supplemental Tables 1-4). Therefore, other imaging sequences may be more sensitive to other histopathological substrates in FTLD spectrum and can be compared to T2*w MRI to further elucidate patterns of degeneration in distinct proteinopathies.

Methodological confounds could also contribute to our findings, but we performed careful histological procedures using optimized methods for iron-stain,^62,63^ run in triplicate and confirmed with comparison to ferritin IHC (Supplemental Figure 3). Finally, given the relatively short fixation interval (*i*.*e*., < 2 years) iron leakage or other tissue degradation is likely to have been minimal in our samples.^91^

While the diversity of our samples’ clinical and pathological features has enabled us to detect unique and frank iron-rich features on joint MRI/histopathology, it is important to acknowledge the limitation of our small sample size in a pathologically heterogeneous disorder. Our findings raise questions regarding the role of inflammation and oxidative stress, but these are dynamic processes with likely varying rates of progression across regions and individual patients. Therefore, we cannot entirely rule out an alternative possibility that all pathological forms of FTLD undergo a similar mechanism of inflammation and iron-overload, captured here at different stages in individual patient samples. Nonetheless, our findings of distinct laminar iron-rich features and corresponding gliosis in areas with high disease burden may also be evident earlier in the disease during life.

Moreover, our work here establishes the basis for future work in larger datasets exploring these focal features. Additionally, our imaging methods, applied in a larger dataset, will enable more quantitative comparison of MRI and digital histology data within and between groups to further inform these complex mechanistic issues.

## 5. CONCLUSION

Our results indicate that FTLD with underlying TDP-43 or tau proteinopathies have distinct mesoscopic signatures within the cortical laminae and neighboring white matter that produce frank features on *ex vivo* T2*w MRI. On histopathology, these features correspond with distinct patterns of microscopic iron-rich pathology, strongly indicative of disparate disease processes amongst FTLD subtypes characterized by prominent upper layer iron-rich astrocytes surrounding small vessels and diffuse iron-rich dystrophic microglia in TDP-43 proteinopathies and hypertrophic microglia in deep layers and adjacent WM in tauopathies. Joint MRI/histopathology studies in FTLD offer a unique method for localizing and studying these focal features — both their spatial distribution across the cortex and their microscopic features.

In addition, previous work has demonstrated that *in vivo* 7T MRI can be sensitized to the intra-cortical pathologic iron content due to AD^33,60,92^ and ALS^37,93^, although not with sufficient resolution to clearly resolve the layer-specific effects we describe here. However, recent work in a small number of healthy subjects has shown the feasibility of increasing *in vivo* cortical laminar resolution of 7T MRI to 250 µm in focal regions, enabling detection of layer-specific myleoarchitecture within small patches of cortex^94^. We therefore expect that the novel iron-rich pathology described here may form a basis for future *in vivo* imaging biomarkers to localize disease processes, distinguish TDP-43 and tau proteinopathies, and further stratify FTLD subtypes.

## ACKNOWLEDGEMENTS

We thank the patients and families for their participation in research for without their important contribution this work would not be possible. We thank Marjolein Bulk for advice in the optimization of iron staining in our tissue. We thank Theresa Schuck and John Robinson for their assistance in tissue harvesting for this project.

## References

1. Rossor MN, Fox NC, Mummery CJ, Schott JM, Warren JD. The diagnosis of young-onset dementia. The Lancet Neurology. 2010;9(8):793–806. doi:10.1016/S1474-4422(10)70159-9

2. Mackenzie IRA, Neumann M, Bigio EH, et al. Nomenclature and nosology for neuropathologic subtypes of frontotemporal lobar degeneration: an update. Acta Neuropathol. 2010;119(1):1. doi:10.1007/s00401-009-0612-2

3. Irwin DJ, Cairns NJ, Grossman M, et al. Frontotemporal lobar degeneration: defining phenotypic diversity through personalized medicine. Acta Neuropathol. 2015;129(4):469–491. doi:10.1007/s00401-014-1380-1

4. Gorno-Tempini ML, Hillis AE, Weintraub S, et al. Classification of primary progressive aphasia and its variants. Neurology. 2011;76(11):1006–1014. doi:10.1212/WNL.0b013e31821103e6

5. Höglinger GU, Respondek G, Stamelou M, et al. Clinical diagnosis of progressive supranuclear palsy: The movement disorder society criteria: MDS Clinical Diagnostic Criteria for PSP. Movement Disorders. 2017;32(6):853–864. doi:10.1002/mds.26987

6. Giannini LAA, Xie SX, McMillan CT, et al. Divergent patterns of TDP□43 and tau pathologies in primary progressive aphasia. Annals of Neurology. 2019;85(5):630–643. doi:10.1002/ana.25465

7. Irwin DJ, McMillan CT, Xie SX, et al. Asymmetry of post-mortem neuropathology in behavioural-variant frontotemporal dementia. Brain. 2018;141(1):288–301. doi:10.1093/brain/awx319

8. Perry DC, Brown JA, Possin KL, et al. Clinicopathological correlations in behavioural variant frontotemporal dementia. Brain. 2017;140(12):3329–3345. doi:10.1093/brain/awx254

9. Spinelli EG, Mandelli ML, Miller ZA, et al. Typical and atypical pathology in primary progressive aphasia variants: Pathology in PPA Variants. Annals of Neurology. 2017;81(3):430–443. doi:10.1002/ana.24885

10. Gordon E, Rohrer JD, Fox NC. Advances in neuroimaging in frontotemporal dementia. Journal of Neurochemistry. 2016;138:193-210. doi:10.1111/jnc.13656

11. Meeter LH, Kaat LD, Rohrer JD, van Swieten JC. Imaging and fluid biomarkers in frontotemporal dementia. Nature Reviews Neurology. 2017;13(7):406–419. doi:10.1038/nrneurol.2017.75

12. Collins JA, Montal V, Hochberg D, et al. Focal temporal pole atrophy and network degeneration in semantic variant primary progressive aphasia. Brain. 2017;140(2):457–471. doi:10.1093/brain/aww313

13. Grossman M, Powers J, Ash S, et al. Disruption of large-scale neural networks in non-fluent/agrammatic variant primary progressive aphasia associated with frontotemporal degeneration pathology. Brain and Language. 2013;127(2):106–120. doi:10.1016/j.bandl.2012.10.005

14. Rohrer JD, Warren JD, Modat M, et al. Patterns of cortical thinning in the language variants of frontotemporal lobar degeneration. Neurology. 2009;72(18):1562–1569.

15. Seeley WW, Crawford RK, Zhou J, Miller BL, Greicius MD. Neurodegenerative Diseases Target Large-Scale Human Brain Networks. Neuron. 2009;62(1):42–52. doi:10.1016/j.neuron.2009.03.024

16. Rohrer JD, Lashley T, Schott JM, et al. Clinical and neuroanatomical signatures of tissue pathology in frontotemporal lobar degeneration. Brain. 2011;134(9):2565–2581. doi:10.1093/brain/awr198

17. Whitwell JL, Jack CR, Parisi JE, et al. Imaging Signatures of Molecular Pathology in Behavioral Variant Frontotemporal Dementia. Journal of Molecular Neuroscience. 2011;45(3). doi:10.1007/s12031-011-9533-3

18. Whitwell JL, Josephs KA. Neuroimaging in frontotemporal lobar degeneration— predicting molecular pathology. Nature Reviews Neurology. 2012;8(3):131–142. doi:10.1038/nrneurol.2012.7

19. Dickson DW, Kouri N, Murray ME, Josephs KA. Neuropathology of Frontotemporal Lobar Degeneration-Tau (FTLD-Tau). Journal of Molecular Neuroscience. 2011;45(3):384–389. doi:10.1007/s12031-011-9589-0

20. Irwin DJ, Brettschneider J, McMillan CT, et al. Deep clinical and neuropathological phenotyping of Pick disease: Deep Phenotype Pick Disease. Annals of Neurology. 2016;79(2):272–287. doi:10.1002/ana.24559

21. Kovacs GG, Lukic MJ, Irwin DJ, et al. Distribution patterns of tau pathology in progressive supranuclear palsy. Acta Neuropathologica. 2020;140(2):99–119. doi:10.1007/s00401-020-02158-2

22. Mackenzie IR, Neumann M. Subcortical TDP-43 pathology patterns validate cortical FTLD-TDP subtypes and demonstrate unique aspects of C9orf72 mutation cases. Acta Neuropathologica. 2020;139(1):83–98. doi:10.1007/s00401-019-02070-4

23. Sakae N, Roemer SF, Bieniek KF, et al. Microglia in frontotemporal lobar degeneration with progranulin or C9ORF72 mutations. Annals of Clinical and Translational Neurology. 2019;6(9):1782–1796. doi:10.1002/acn3.50875

24. Bulk M, Abdelmoula WM, Geut H, et al. Quantitative MRI and laser ablation-inductively coupled plasma-mass spectrometry imaging of iron in the frontal cortex of healthy controls and Alzheimer’s disease patients. NeuroImage. 2020;215:116808. doi:10.1016/j.neuroimage.2020.116808

25. Fracasso A, van Veluw SJ, Visser F, et al. Lines of Baillarger in vivo and ex vivo: Myelin contrast across lamina at 7 T MRI and histology. NeuroImage. 2016;133:163-175. doi:10.1016/j.neuroimage.2016.02.072

26. Fukunaga M, Li T-Q, van Gelderen P, et al. Layer-specific variation of iron content in cerebral cortex as a source of MRI contrast. Proceedings of the National Academy of Sciences. 2010;107(8):3834–3839. doi:10.1073/pnas.0911177107

27. Hametner S, Endmayr V, Deistung A, et al. The influence of brain iron and myelin on magnetic susceptibility and effective transverse relaxation -A biochemical and histological validation study. NeuroImage. 2018;179:117-133. doi:10.1016/j.neuroimage.2018.06.007

28. Wallace MN, Cronin MJ, Bowtell RW, Scott IS, Palmer AR, Gowland PA. Histological Basis of Laminar MRI Patterns in High Resolution Images of Fixed Human Auditory Cortex. Frontiers in Neuroscience. 2016;10. doi:10.3389/fnins.2016.00455

29. Cheli VT, Correale J, Paez PM, Pasquini JM. Iron Metabolism in Oligodendrocytes and Astrocytes, Implications for Myelination and Remyelination. ASN Neuro. 2020;12:175909142096268. doi:10.1177/1759091420962681

30. Benveniste H, Einstein G, Kim KR, Hulette C, Johnson GA. Detection of neuritic plaques in Alzheimer’s disease by magnetic resonance microscopy. Proceedings of the National Academy of Sciences. 1999;96(24):14079–14084. doi:10.1073/pnas.96.24.14079

31. Bulk M, Abdelmoula WM, Nabuurs RJA, et al. Postmortem MRI and histology demonstrate differential iron accumulation and cortical myelin organization in early- and late-onset Alzheimer’s disease. Neurobiology of Aging. 2018;62:231-242. doi:10.1016/j.neurobiolaging.2017.10.017

32. Bulk M, Kenkhuis B, van der Graaf LM, Goeman JJ, Natté R, van der Weerd L. Postmortem T2*-Weighted MRI Imaging of Cortical Iron Reflects Severity of Alzheimer’s Disease. Bush A, ed. Journal of Alzheimer’s Disease. 2018;65(4):1125–1137. doi:10.3233/JAD-180317

33. Gong N-J, Dibb R, Bulk M, van der Weerd L, Liu C. Imaging beta amyloid aggregation and iron accumulation in Alzheimer’s disease using quantitative susceptibility mapping MRI. NeuroImage. 2019;191:176–185. doi:10.1016/j.neuroimage.2019.02.019

34. Meadowcroft MD, Connor JR, Smith MB, Yang QX. MRI and histological analysis of beta-amyloid plaques in both human Alzheimer’s disease and APP/PS1 transgenic mice. Journal of Magnetic Resonance Imaging. 2009;29(5):997–1007. doi:10.1002/jmri.21731

35. van Rooden S, Maat-Schieman MLC, Nabuurs RJA, et al. Cerebral Amyloidosis: Postmortem Detection with Human 7.0-T MR Imaging System. Radiology. 2009;253(3):788–796. doi:10.1148/radiol.2533090490

36. Zeineh MM, Chen Y, Kitzler HH, Hammond R, Vogel H, Rutt BK. Activated iron-containing microglia in the human hippocampus identified by magnetic resonance imaging in Alzheimer disease. Neurobiology of Aging. 2015;36(9):2483–2500. doi:10.1016/j.neurobiolaging.2015.05.022

37. Kwan JY, Jeong SY, Van Gelderen P, et al. Iron Accumulation in Deep Cortical Layers Accounts for MRI Signal Abnormalities in ALS: Correlating 7 Tesla MRI and Pathology. Ashizawa T, ed. PLoS ONE. 2012;7(4):e35241. doi:10.1371/journal.pone.0035241

38. Meadowcroft MD, Mutic NJ, Bigler DC, et al. Histological-MRI correlation in the primary motor cortex of patients with amyotrophic lateral sclerosis: MRI and Histological Analysis of the PMC in ALS. Journal of Magnetic Resonance Imaging. 2015;41(3):665–675. doi:10.1002/jmri.24582

39. Pallebage-Gamarallage M, Foxley S, Menke RAL, et al. Dissecting the pathobiology of altered MRI signal in amyotrophic lateral sclerosis: A post mortem whole brain sampling strategy for the integration of ultra-high-field MRI and quantitative neuropathology. BMC Neuroscience. 2018;19(1). doi:10.1186/s12868-018-0416-1

40. Wang C, Foxley S, Ansorge O, et al. Methods for quantitative susceptibility and R2* mapping in whole post-mortem brains at 7T applied to amyotrophic lateral sclerosis. NeuroImage. 2020;222:117216. doi:10.1016/j.neuroimage.2020.117216

41. De Reuck JL, Deramecourt V, Auger F, et al. Iron deposits in post-mortem brains of patients with neurodegenerative and cerebrovascular diseases: a semi-quantitative 7.0 T magnetic resonance imaging study. European Journal of Neurology. 2014;21(7):1026–1031. doi:10.1111/ene.12432

42. Foroutan P, Murray ME, Fujioka S, et al. Progressive Supranuclear Palsy: High-Field-Strength MR Microscopy in the Human Substantia Nigra and Globus Pallidus. Radiology. 2013;266(1):280–288. doi:10.1148/radiol.12102273

43. Massey L, Miranda M, Al-Helli O, et al. 9.4 T MR microscopy of the substantia nigra with pathological validation in controls and disease. NeuroImage: Clinical. 2017;13:154–163. doi:10.1016/j.nicl.2016.11.015

44. Armstrong RA, Cairns NJ, Lantos PL. Laminar distribution of Pick bodies, Pick cells and Alzheimer disease pathology in the frontal and temporal cortex in Pick’s disease. Neuropathology and Applied Neurobiology. 1999;25(4):266–271. doi:10.1046/j.1365-2990.1999.00173.x

45. Armstrong RA, Cairns NJ. Different molecular pathologies result in similar spatial patterns of cellular inclusions in neurodegenerative disease: a comparative study of eight disorders. J Neural Transm. 2012;119(12):1551–1560. doi:10.1007/s00702-012-0838-3

46. Kersaitis C, Halliday GM, Kril JJ. Regional and cellular pathology in frontotemporal dementia: relationship to stage of disease in cases with and without Pick bodies. Acta Neuropathologica. 2004;108(6):515–523. doi:10.1007/s00401-004-0917-0

47. Schofield E. Severity of gliosis in Pick’s disease and frontotemporal lobar degeneration: tau-positive glia differentiate these disorders. Brain. 2003;126(4):827–840. doi:10.1093/brain/awg085

48. Hasegawa I, Takeda A, Hatsuta H, et al. An autopsy case of globular glial tauopathy presenting with clinical features of motor neuron disease with dementia and iron deposition in the motor cortex: Globular glial tauopathy. Neuropathology. 2018;38(4):372–379. doi:10.1111/neup.12457

49. Cooper PN, Siddons CA, Mann DMA. Patterns of glial cell activity in fronto-temporal dementia (lobar atrophy). Neuropathology and Applied Neurobiology. 1996;22(1):17–22. doi:10.1111/j.1365-2990.1996.tb00841.x

50. McMillan CT, Irwin DJ, Avants BB, et al. White matter imaging helps dissociate tau from TDP-43 in frontotemporal lobar degeneration. Journal of Neurology, Neurosurgery & Psychiatry. 2013;84(9):949–955. doi:10.1136/jnnp-2012-304418

51. Giannini LAA, Peterson C, Ohm D, et al. Frontotemporal lobar degeneration proteinopathies have disparate microscopic patterns of white and grey matter pathology. acta neuropathol commun. 2021;9(1):30. doi:10.1186/s40478-021-01129-2

52. Armstrong RA, Hamilton RL, Mackenzie IRA, Hedreen J, Cairns NJ. Laminar distribution of the pathological changes in sporadic frontotemporal lobar degeneration with transactive response (TAR) DNA-binding protein of 43 kDa (TDP-43) proteinopathy: a quantitative study using polynomial curve fitting: Laminar pathology in sporadic FTLD-TDP. Neuropathology and Applied Neurobiology. 2013;39(4):335–347. doi:10.1111/j.1365-2990.2012.01291.x

53. Armstrong RA, Carter D, Cairns NJ. A quantitative study of the neuropathology of 32 sporadic and familial cases of frontotemporal lobar degeneration with TDP-43 proteinopathy (FTLD-TDP): Familial and sporadic FTLD-TDP. Neuropathology and Applied Neurobiology. 2012;38(1):25–38. doi:10.1111/j.1365-2990.2011.01188.x

54. Lant SB, Robinson AC, Thompson JC, et al. Patterns of microglial cell activation in frontotemporal lobar degeneration: Microglia and frontotemporal lobar degeneration. Neuropathology and Applied Neurobiology. 2014;40(6):686–696. doi:10.1111/nan.12092

55. Taipa R, Brochado P, Robinson A, et al. Patterns of Microglial Cell Activation in Alzheimer Disease and Frontotemporal Lobar Degeneration. Neurodegenerative Diseases. 2017;17(4-5):145–154. doi:10.1159/000457127

56. Rascovsky K, Hodges JR, Knopman D, et al. Sensitivity of revised diagnostic criteria for the behavioural variant of frontotemporal dementia. Brain. 2011;134(9):2456–2477. doi:10.1093/brain/awr179

57. Toledo JB, Van Deerlin VM, Lee EB, et al. A platform for discovery: The University of Pennsylvania Integrated Neurodegenerative Disease Biobank. Alzheimer’s & Dementia. 2014;10(4):477-484.e1. doi:10.1016/j.jalz.2013.06.003

58. Mackenzie IRA, Neumann M, Baborie A, et al. A harmonized classification system for FTLD-TDP pathology. Acta Neuropathologica. 2011;122(1):111–113. doi:10.1007/s00401-011-0845-8

59. Montine TJ, Phelps CH, Beach TG, et al. National Institute on Aging–Alzheimer’s Association guidelines for the neuropathologic assessment of Alzheimer’s disease: a practical approach. Acta Neuropathologica. 2012;123(1):1–11. doi:10.1007/s00401-011-0910-3

60. Kenkhuis B, Jonkman LE, Bulk M, et al. 7T MRI allows detection of disturbed cortical lamination of the medial temporal lobe in patients with Alzheimer’s disease. NeuroImage: Clinical. 2019;21:101665.

61. Irwin DJ, Byrne MD, McMillan CT, et al. Semi-Automated Digital Image Analysis of Pick’s Disease and TDP-43 Proteinopathy. Journal of Histochemistry & Cytochemistry. 2016;64(1):54–66. doi:10.1369/0022155415614303

62. Meguro R, Asano Y, Odagiri S, Li C, Iwatsuki H, Shoumura K. Nonheme-iron histochemistry for light and electron microscopy: a historical, theoretical and technical review. Archives of histology and cytology. 2007;70(1):1–19.

63. van Duijn S, Nabuurs RJA, van Duinen SG, Natté R. Comparison of Histological Techniques to Visualize Iron in Paraffin-embedded Brain Tissue of Patients with Alzheimer’s Disease. Journal of Histochemistry & Cytochemistry. 2013;61(11):785–792. doi:10.1369/0022155413501325

64. Lee EB, Skovronsky DM, Abtahian F, Doms RW, Lee VM-Y. Secretion and Intracellular Generation of Truncated Aβ in β-Site Amyloid-β Precursor Protein-cleaving Enzyme Expressing Human Neurons. Journal of Biological Chemistry. 2003;278(7):4458–4466. doi:10.1074/jbc.M210105200

65. Mercken M, Vandermeeren M, Lübke U, et al. Monoclonal antibodies with selective specificity for Alzheimer Tau are directed against phosphatase-sensitive epitopes. Acta Neuropathologica. 1992;84(3):265–272. doi:10.1007/BF00227819

66. Neumann M, Kwong LK, Lee EB, et al. Phosphorylation of S409/410 of TDP-43 is a consistent feature in all sporadic and familial forms of TDP-43 proteinopathies. Acta Neuropathologica. 2009;117(2):137–149. doi:10.1007/s00401-008-0477-9

67. Bachstetter AD, Van Eldik LJ, Schmitt FA, et al. Disease-related microglia heterogeneity in the hippocampus of Alzheimer’s disease, dementia with Lewy bodies, and hippocampal sclerosis of aging. Acta Neuropathologica Communications. 2015;3(1). doi:10.1186/s40478-015-0209-z

68. Ito D, Imai Y, Ohsawa K, Nakajima K, Fukuuchi Y, Kohsaka S. Microglia-specific localisation of a novel calcium binding protein, Iba1. Molecular Brain Research. 1998;57(1):1–9. doi:10.1016/S0169-328X(98)00040-0

69. Beach TG, Walker R, McGeer EG. Patterns of gliosis in alzheimer’s disease and aging cerebrum. Glia. 1989;2(6):420–436. doi:10.1002/glia.440020605

70. Lundgaard I, Osório MJ, Kress BT, Sanggaard S, Nedergaard M. White matter astrocytes in health and disease. Neuroscience. 2014;276:161–173. doi:10.1016/j.neuroscience.2013.10.050

71. Ahmed Z, Bigio EH, Budka H, et al. Globular glial tauopathies (GGT): consensus recommendations. Acta Neuropathologica. 2013;126(4):537–544. doi:10.1007/s00401-013-1171-0

72. Oba H, Araki T, Ohtomo K, et al. Amyotrophic lateral sclerosis: T2 shortening in motor cortex at MR imaging. Radiology. 1993;189(3):843–846.

73. Brettschneider J, Del Tredici K, Toledo JB, et al. Stages of pTDP-43 pathology in amyotrophic lateral sclerosis: ALS Stages. Annals of Neurology. 2013;74(1):20–38. doi:10.1002/ana.23937

74. Josephs KA, Whitwell JL, Knopman DS, et al. Two distinct subtypes of right temporal variant frontotemporal dementia. Neurology. 2009;73(18):1443–1450. doi:10.1212/WNL.0b013e3181bf9945

75. Biasiotto G, Di Lorenzo D, Archetti S, Zanella I. Iron and Neurodegeneration: Is Ferritinophagy the Link? Molecular Neurobiology. 2016;53(8):5542–5574. doi:10.1007/s12035-015-9473-y

76. Rao SS, Adlard PA. Untangling Tau and Iron: Exploring the Interaction Between Iron and Tau in Neurodegeneration. Frontiers in Molecular Neuroscience. 2018;11. doi:10.3389/fnmol.2018.00276

77. Ward RJ, Zucca FA, Duyn JH, Crichton RR, Zecca L. The role of iron in brain ageing and neurodegenerative disorders. The Lancet Neurology. 2014;13(10):1045–1060. doi:10.1016/S1474-4422(14)70117-6

78. Heppner FL, Ransohoff RM, Becher B. Immune attack: the role of inflammation in Alzheimer disease. Nature Reviews Neuroscience. 2015;16(6):358–372. doi:10.1038/nrn3880

79. Ransohoff RM. How neuroinflammation contributes to neurodegeneration. Science. 2016;353(6301):777–783. doi:10.1126/science.aag2590

80. Boche D, Perry VH, Nicoll JAR. Review: Activation patterns of microglia and their identification in the human brain: Microglia in human brain. Neuropathology and Applied Neurobiology. 2013;39(1):3–18. doi:10.1111/nan.12011

81. Ishizawa K, Dickson DW. Microglial Activation parallels System Degeneration in progressive Supranuclear palsy and Corticobasal Degeneration. Journal of Neuropathology & Experimental Neurology. 2001;60(6):647–657. doi:10.1093/jnen/60.6.647

82. Bellucci A, Bugiani O, Ghetti B, Spillantini MG. Presence of Reactive Microglia and Neuroinflammatory Mediators in a Case of Frontotemporal Dementia with P301S Mutation. Neurodegenerative Diseases. 2011;8(4):221–229. doi:10.1159/000322228

83. Streit WJ, Khoshbouei H, Bechmann I. Dystrophic microglia in late□onset Alzheimer’s disease. Glia. 2020;68(4):845–854. doi:10.1002/glia.23782

84. Streit WJ, Sammons NW, Kuhns AJ, Sparks DL. Dystrophic microglia in the aging human brain. Glia. 2004;45(2):208–212. doi:10.1002/glia.10319

85. Ohm DT, Kim G, Gefen T, et al. Prominent microglial activation in cortical white matter is selectively associated with cortical atrophy in primary progressive aphasia. Neuropathology and Applied Neurobiology. 2019;45(3):216–229. doi:10.1111/nan.12494

86. Hsu ET, Gangolli M, Su S, et al. Astrocytic degeneration in chronic traumatic encephalopathy. Acta Neuropathologica. 2018;136(6):955–972. doi:10.1007/s00401-018-1902-3

87. Martinac JA, Craft DK, Su JH, Kim RC, Cotman CW. Astrocytes degenerate in frontotemporal dementia: possible relation to hypoperfusion. Neurobiology of Aging. 2001;22(2):195–207. doi:10.1016/S0197-4580(00)00231-1

88. Lee EB, Porta S, Michael Baer G, et al. Expansion of the classification of FTLD-TDP: distinct pathology associated with rapidly progressive frontotemporal degeneration. Acta Neuropathologica. 2017;134(1):65–78. doi:10.1007/s00401-017-1679-9

89. Pelizzoni I, Zacchetti D, Campanella A, Grohovaz F, Codazzi F. Iron uptake in quiescent and inflammation-activated astrocytes: A potentially neuroprotective control of iron burden. Biochimica et Biophysica Acta (BBA) -Molecular Basis of Disease. 2013;1832(8):1326–1333. doi:10.1016/j.bbadis.2013.04.007

90. van Duijn S, Bulk M, van Duinen SG, et al. Cortical Iron Reflects Severity of Alzheimer’s Disease. Journal of Alzheimer’s Disease. 2017;60(4):1533–1545. doi:10.3233/JAD-161143

91. Gellein K, Flaten TP, Erikson KM, Aschner M, Syversen T. Leaching of Trace Elements from Biological Tissue by Formalin Fixation. Biological Trace Element Research. 2008;121(3):221–225. doi:10.1007/s12011-007-8051-1

92. van Rooden S, Versluis MJ, Liem MK, et al. Cortical phase changes in Alzheimer’s disease at 7T MRI: A novel imaging marker. Alzheimer’s & Dementia. 2014;10(1):e19–e26. doi:10.1016/j.jalz.2013.02.002

93. Acosta-Cabronero J, Machts J, Schreiber S, et al. Quantitative Susceptibility MRI to Detect Brain Iron in Amyotrophic Lateral Sclerosis. Radiology. 2018;289(1):195–203. doi:10.1148/radiol.2018180112

94. Balasubramanian M, Mulkern RV, Neil JJ, Maier SE, Polimeni JR. Probing in vivo cortical myeloarchitecture in humans via line-scan diffusion acquisitions at 7 T with 250-500 micron radial resolution. Magnetic Resonance in Medicine. Published online August 1, 2020. doi:10.1002/mrm.28419

