## Supplementary Materials for "Joint *ex vivo* MRI and histology detect iron-rich cortical gliosis in Tau and TDP-43 proteinopathies"

### Supplementary Table 1. Neuropathological Ratings.

| Patient | Region | Layer | NL | Inclusion | Amyloid | Fe | Ferritin | IBA1 | GFAP |
|---|---|---|---|---|---|---|---|---|---|
| 1 (HC) | OFC | Upper GM | 0 | 0 | 0 | 0 | 0.5 | 1 | 1 |
| | | Deep GM | 0 | 0 | 0 | 0 | 0.5 | 1 | 0 |
| | | WM-U | N/A | 0 | 0 | 0 | 0 | 1 | 3 |
| | | WM Deep | N/A | 0 | 0 | 0 | 0 | 1 | 3 |
| | lPFC | Upper GM | 0 | 0 | 0 | 0.5 | 0.5 | 0.5 | 1 |
| | | Deep GM | 0 | 0 | 0 | 0.5 | 0.5 | 0.5 | 0 |
| | | WM-U | N/A | 0 | 0 | 0 | 0 | 1 | 3 |
| | | WM Deep | N/A | 0 | 0 | 0 | 0 | 1 | 3 |
| | MOT | Upper GM | 0 | 0 | 0.5 | 0 | 0.5 | 0.5 | 0 |
| | | Deep GM | 0 | 0 | 0 | 0 | 0.5 | 0.5 | 0 |
| | | WM-U | N/A | 0 | 0 | 0 | 0.5 | 0.5 | 3 |
| | | WM Deep | N/A | 0 | 0 | 0 | 0.5 | 0.5 | 3 |
| | SENS | Upper GM | 0 | 0 | 0.5 | 0 | 0.5 | 0.5 | 1 |
| | | Deep GM | 0 | 0 | 0.5 | 0 | 0.5 | 0.5 | 0 |
| | | WM-U | N/A | 0 | 0 | 0 | 0 | 0.5 | 3 |
| | | WM Deep | N/A | 0 | 0 | 0 | 0 | 0.5 | 3 |
| 2 (AD) | ATP | Upper GM | 2 | 1 | 3 | 2 | 3 | 3 | 3 |
| | | Deep GM | 2 | 1 | 3 | 3 | 3 | 3 | 2 |
| | | WM-U | N/A | | 0.5 | 2 | 2 | 1 | 3 |
| | | WM Deep | N/A | | 0.5 | 1 | 2 | 1 | 2 |
| | OFC | Upper GM | 1 | 3 | 3 | 2 | 2 | 2 | 2 |
| | | Deep GM | 1 | 3 | 3 | 3 | 2 | 2 | 2 |
| | | WM-U | N/A | 0 | 0 | 1 | 0.5 | 2 | 2 |
| | | WM Deep | N/A | 0 | 0 | 0 | 0.5 | 2 | 2 |
| | lPFC | Upper GM | 2 | 3 | 3 | 1 | 2 | 1 | 2 |
| | | Deep GM | 1 | 3 | 3 | 2 | 2 | 2 | 2 |
| | | WM-U | N/A | 0.5 | 0.5 | 0.5 | 2 | 1 | 2 |
| | | WM Deep | N/A | 0.5 | 0.5 | 0 | 2 | 1 | 2 |
| | MOT | Upper GM | 0 | 2 | 2 | 1 | 2 | 1 | 1 |
| | | Deep GM | 0 | 2 | 2 | 2 | 2 | 1 | 1 |
| | | WM-U | N/A | 0.5 | 0 | 1 | 0 | 1 | 2 |
| | | WM Deep | N/A | 0.5 | 0 | 0 | 0 | 1 | 2 |
| | SENS | Upper GM | 1 | 3 | 3 | 0 | 2 | 2 | 2 |
| | | Deep GM | 1 | 3 | 3 | 1 | 2 | 2 | 2 |
| | | WM-U | N/A | 0 | 0 | 0 | 1 | 2 | 2 |
| | | WM Deep | N/A | 0 | 0 | 0 | 1 | 2 | 2 |
| 3(TDPC) | ATP | Upper GM | 3 | 3 | 0 | 3 | 2 | 3 | 3 |
| | | Deep GM | 2 | 3 | 0 | 2 | 1 | 3 | 3 |
| | | WM-U | N/A | 0.5 | 0 | 1 | 1 | 3 | 3 |
| | | WM Deep | N/A | 0.5 | 0 | 1 | 1 | 3 | 3 |
| | OFC | Upper GM | 2 | 2 | 0 | 1 | 2 | 3 | 2 |
| | | Deep GM | 1 | 1 | 0 | 1 | 2 | 3 | 0 |
| | | WM-U | N/A | 0.5 | 0 | 0.5 | 0.5 | 3 | 2 |
| | | WM Deep | N/A | 0.5 | 0 | 0.5 | 0.5 | 3 | 1 |
| | lPFC | Upper GM | 1 | 1 | 0 | 0.5 | 1 | 1 | 1 |
| | | Deep GM | 0 | 0.5 | 0.5 | 0.5 | 1 | 0.5 | 0 |
| | | WM-U | N/A | 0 | 0 | 0 | 1 | 1 | 2 |
| | | WM Deep | N/A | 0 | 0 | 0 | 1 | 0.5 | 2 |
| | MOT | Upper GM | 0 | 0.5 | 0 | 0.5 | 1 | 1 | 1 |
| | | Deep GM | 0 | 0 | 0 | 0 | 1 | 1 | 0 |
| | | WM-U | N/A | 0 | 0 | 0 | 1 | 1 | 2 |
| | | WM Deep | N/A | 0 | 0 | 0 | 1 | 1 | 2 |
| | SENS | Upper GM | 0 | 0 | 0 | 0.5 | 1 | 1 | 1 |

| | | | | | | | | | |
|---|---|---|---|---|---|---|---|---|---|
| | | Deep GM | 0 | 0 | 0 | 0.5 | 1 | 1 | 0 |
| | | WM-U | N/A | 0 | 0 | 0 | 0.5 | 1 | 2 |
| | | WM Deep | N/A | 0 | 0 | 0 | 0.5 | 1 | 2 |
| 4(TDPA) | ATP | Upper GM | 2 | 2 | 0 | 1 | 3 | 3 | 2 |
| | | Deep GM | 1 | 1 | 0 | 2 | 2 | 2 | 1 |
| | | WM-U | N/A | 0.5 | 0 | 1 | 1 | 1 | 2 |
| | | WM Deep | N/A | 0.5 | 0 | 1 | 1 | 1 | 1 |
| | OFC | Upper GM | 3 | 3 | 0 | 2 | 3 | 3 | 3 |
| | | Deep GM | 1 | 0.5 | 0 | 2 | 2 | 2 | 2 |
| | | WM-U | N/A | 0.5 | 0 | 1 | 1 | 1 | 2 |
| | | WM Deep | N/A | 0.5 | 0 | 1 | 1 | 1 | 0.5 |
| | lOFC (MRI) | Upper GM | 2 | 3 | 0 | 3 | 3 | 2 | 3 |
| | | Deep GM | 0 | 1 | 0 | 2 | 2 | 1 | 2 |
| | | WM-U | N/A | 1 | 0 | 1 | 1 | 0.5 | 3 |
| | | WM Deep | N/A | 1 | 0 | 1 | 1 | 0.5 | 3 |
| | IPFC | Upper GM | 1 | 1 | 0 | 0.5 | 2 | 1 | 1 |
| | | Deep GM | 0 | 0 | 0 | 0 | 1 | 0.5 | 0 |
| | | WM-U | N/A | 0 | 0 | 0 | 1 | 0.5 | 2 |
| | | WM Deep | N/A | 0 | 0 | 0 | 1 | 0.5 | 2 |
| | MOT | Upper GM | 1 | 1 | 0 | 1 | 2 | 2 | 1 |
| | | Deep GM | 0 | 0 | 0 | 1 | 1 | 1 | 0 |
| | | WM-U | N/A | 0 | 0 | 0 | 1 | 1 | 2 |
| | | WM Deep | N/A | 0 | 0 | 0 | 1 | 1 | 2 |
| | SENS | Upper GM | 0 | 0 | 0 | 0.5 | 1 | 1 | 0.5 |
| | | Deep GM | 0 | 0 | 0 | 0.5 | 0.5 | 1 | 0 |
| | | WM-U | N/A | 0 | 0 | 0 | 0 | 1 | 2 |
| | | WM Deep | N/A | 0 | 0 | 0 | 0 | 1 | 2 |
| 5 (GGT) | OFC | Upper GM | 0 | 0.5 | 2 | 0 | 1 | 1 | 1 |
| | | Deep GM | 0 | 0.5 | 2 | 0.5 | 1 | 1 | 1 |
| | | WM-U | N/A | 0.5 | 0 | 0 | 1 | 2 | 2 |
| | | WM Deep | N/A | 0.5 | 0 | 0 | 1 | 2 | 2 |
| | iMOT(MRI) | Upper GM | 3 | 3 | 0.5 | 1 | 3 | 3 | 3 |
| | | Deep GM | 3 | 3 | 0.5 | 3 | 3 | 3 | 3 |
| | | WM-U | N/A | 3 | 0 | 2 | 3 | 3 | 1 |
| | | WM Deep | N/A | 3 | 0 | 2 | 3 | 3 | 1 |
| | IPFC | Upper GM | 1 | 1 | 2 | 1 | 2 | 2 | 2 |
| | | Deep GM | 1 | 1 | 2 | 2 | 2 | 2 | 2 |
| | | WM-U | N/A | 1 | 0 | 0.5 | 1 | 3 | 2 |
| | | WM Deep | N/A | 1 | 0.5 | 0.5 | 1 | 3 | 2 |
| | MOT | Upper GM | 3 | 2 | 0.5 | 1 | 3 | 3 | 3 |
| | | Deep GM | 3 | 3 | 0.5 | 3 | 3 | 3 | 3 |
| | | WM-U | N/A | 3 | 0 | 2 | 3 | 3 | 2 |
| | | WM Deep | N/A | 3 | 0 | 2 | 3 | 3 | 1 |
| | SENS | Upper GM | 0 | 0.5 | 0.5 | 0 | 2 | 2 | 2 |
| | | Deep GM | 0 | 0.5 | 0.5 | 0 | 2 | 2 | 2 |
| | | WM-U | N/A | 1 | 0 | 0 | 2 | 3 | 2 |
| | | WM Deep | N/A | 1 | 0 | 0 | 2 | 3 | 2 |
| 6 (PSP) | ATP | Upper GM | 0 | 1 | 2 | 2 | 2 | 2 | 2 |
| | | Deep GM | 0 | 1 | 2 | 2 | 2 | 2 | 1 |
| | | WM-U | N/A | 0.5 | 0 | 0.5 | 2 | 3 | 2 |
| | | WM Deep | N/A | 0.5 | 0 | 0.5 | 2 | 3 | 2 |
| | OFC | Upper GM | 0 | 0.5 | 3 | 0.5 | 3 | 2 | 1 |
| | | Deep GM | 0 | 0.5 | 3 | 1 | 3 | 2 | 0 |
| | | WM-U | N/A | 0.5 | 0 | 1 | 2 | 3 | 2 |
| | | WM Deep | N/A | 0.5 | 0 | 1 | 2 | 3 | 2 |
| | IPFC | Upper GM | 1 | 1 | 2 | 1 | 3 | 2 | 2 |
| | | Deep GM | 1 | 2 | 2 | 2 | 3 | 2 | 1 |
| | | WM-U | N/A | 1 | 0 | 1 | 2 | 3 | 2 |
| | | WM Deep | N/A | 1 | 0 | 1 | 2 | 3 | 2 |
| | MOT | Upper GM | 1 | 2 | 0.5 | 1 | 3 | 3 | 3 |
| | | Deep GM | 1 | 3 | 0.5 | 3 | 3 | 2 | 3 |
| | | WM-U | N/A | 2 | 0 | 2 | 2 | 3 | 3 |
| | | WM Deep | N/A | 2 | 0 | 2 | 2 | 3 | 3 |
| | SENS | Upper GM | 0 | 1 | 1 | 0.5 | 2 | 1 | 1 |
| | | Deep GM | 0 | 1 | 1 | 1 | 2 | 1 | 1 |
| | | WM-U | N/A | 1 | 0 | 0 | 2 | 1 | 2 |

| | | | | | | | | | |
|---|---|---|---|---|---|---|---|---|---|
| | | WM Deep | N/A | 1 | 0 | 0 | 2 | 2 | 2 |
| 7(PiD) | OFC | Upper GM | 3 | 3 | 0 | 0.5 | 2 | 1 | 2 |
| | | Deep GM | 3 | 3 | 0 | 1 | 2 | 1 | 1 |
| | | WM-U | N/A | 1 | 0 | 2 | 1 | 1 | 2 |
| | | WM Deep | N/A | 1 | 0 | 1 | 1 | 1 | 2 |
| | DLPFC(MRI) | Upper GM | 3 | 3 | 0 | 0.5 | 3 | 3 | 3 |
| | | Deep GM | 3 | 3 | 0 | 2 | 3 | 3 | 3 |
| | | WM-U | N/A | 3 | 0 | 2 | 2 | 1 | 3 |
| | | WM Deep | N/A | 3 | 0 | 1 | 2 | 1 | 3 |
| | IPFC | Upper GM | 3 | 3 | 0 | 0 | 2 | 1 | 2 |
| | | Deep GM | 3 | 3 | 0 | 2 | 2 | 2 | 1 |
| | | WM-U | N/A | 1 | 0 | 2 | 1 | 1 | 2 |
| | | WM Deep | N/A | 1 | 0 | 1 | 1 | 1 | 2 |
| | MOT | Upper GM | 0 | 0.5 | 0 | 0 | 2 | 1 | 0 |
| | | Deep GM | 0 | 0.5 | 0 | 0.5 | 1 | 1 | 0 |
| | | WM-U | N/A | 0 | 0 | 0 | 1 | 1 | 1 |
| | | WM Deep | N/A | 0 | 0 | 0 | 1 | 1 | 1 |
| | SENS | Upper GM | 0 | 0.5 | 0 | 0 | 0.5 | 0.5 | 0.5 |
| | | Deep GM | 0 | 0.5 | 0.5 | 0.5 | 0.5 | 0.5 | 0 |
| | | WM-U | N/A | 0.5 | 0 | 0 | 0 | 1 | 1 |
| | | WM Deep | N/A | 0.5 | 0 | 0 | 0 | 1 | 1 |

Table depicts ordinal ratings in each patient sample region on standard ordinal rating scale (0=none, 0.5=rare, 1=mild, 2=moderate, 3=severe). NL= neuronal loss from LFB stained slides, Inclusion= proteinopathy severity (tau for FTLD-Tau, AD and HC; TDP-43 for FTLD-TDP), Amyloid= amyloid plaque score, Fe= score for glial pathology detected by Perl's stain for iron with DAB enhancement, Ferritin= = score for glial pathology detected by ferritin immunostain, IBA-1= activated microglial score from IBA-1 immunostain, GFAP= activated astrocyte score for GFAP immunostain, HC= healthy control, AD= Alzheimer's disease, TDPA= FTLD-TDP subtype A, TDPC= FTLD-TDP subtype C, GGT= FTLD-Tau with globular glial tauopathy, PSP= FTLD-Tau with progressive supranuclear palsy, PiD= FTLD-Tau with Pick's disease, ATP= anterior temporal pole, OFC= orbitofrontal cortex, IPFC= inferior prefrontal cortex, MOT= primary motor cortex, SENS= primary sensory cortex, lOFC (MRI)= MRI guided sample from lateral OFC, iMOT (MRI)= MRI guided inferior primary motor cortex, DLPFC (MRI)= MRI guided sample from dorsolateral frontal cortex, upper GM= grey matter in upper cortical layers II-III, deep GM= GM in deep cortical layers IV-VI , WM-U= juxtacortical white matter enriched for U-fibers, WM deep= relative deep WM, N/A= not applicable as no neuron loss to assess in WM layers.

### Supplementary Table 2. Histopathological ratings of white matter integrity.

| Patient | Region | Layer | WM LFB | WM Fe |
|---|---|---|---|---|
| 1 (HC) | OFC | Deep GM | 2 | 2 |
| | | WM-U | 3 | 3 |
| | | WM Deep | 3 | 3 |
| | lPFC | Deep GM | 2 | 2 |
| | | WM-U | 3 | 3 |
| | | WM Deep | 3 | 3 |
| | MOT | Deep GM | 3 | 3 |
| | | WM-U | 3 | 3 |
| | | WM Deep | 3 | 3 |
| | SENS | Deep GM | 3 | 3 |
| | | WM-U | 3 | 3 |
| | | WM Deep | 3 | 3 |
| 2 (AD) | ATP | Deep GM | 0.5 | 0.5 |
| | | WM-U | 3 | 3 |
| | | WM Deep | 3 | 3 |
| | OFC | Deep GM | 1 | 1 |
| | | WM-U | 3 | 3 |
| | | WM Deep | 3 | 3 |
| | lPFC | Deep GM | 2 | 2 |
| | | WM-U | 3 | 3 |
| | | WM Deep | 3 | 3 |
| | MOT | Deep GM | 3 | 2 |
| | | WM-U | 3 | 2 |
| | | WM Deep | 3 | 2 |
| | SENS | Deep GM | 3 | 2 |
| | | WM-U | 3 | 3 |
| | | WM Deep | 3 | 3 |
| 3(TDPC) | ATP | Deep GM | 0.5 | 0.5 |
| | | WM-U | 1 | 3 |
| | | WM Deep | 3 | 3 |
| | OFC | Deep GM | 1 | 0.5 |
| | | WM-U | 3 | 3 |
| | | WM Deep | 3 | 3 |
| | lPFC | Deep GM | 2 | 2 |
| | | WM-U | 3 | 3 |
| | | WM Deep | 3 | 3 |
| | MOT | Deep GM | 3 | 3 |
| | | WM-U | 3 | 3 |
| | | WM Deep | 3 | 3 |
| | SENS | Deep GM | 3 | 3 |
| | | WM-U | 3 | 3 |
| | | WM Deep | 3 | 3 |
| 4(TDPA) | ATP | Deep GM | 0.5 | 0.5 |
| | | WM-U | 2 | 3 |
| | | WM Deep | 3 | 3 |
| | OFC | Deep GM | 0.5 | 0.5 |
| | | WM-U | 2 | 2 |
| | | WM Deep | 1 | 1 |
| | lOFC (MRI) | Deep GM | 1 | 1 |
| | | WM-U | 2 | 3 |
| | | WM Deep | 2 | 3 |
| | lPFC | Deep GM | 2 | 2 |
| | | WM-U | 3 | 2 |
| | | WM Deep | 3 | 2 |
| | MOT | Deep GM | 3 | 3 |
| | | WM-U | 3 | 2 |
| | | WM Deep | 3 | 2 |
| | SENS | Deep GM | 3 | 2 |
| | | WM-U | 3 | 2 |
| | | WM Deep | 3 | 2 |
| 5 (GGT) | OFC | Deep GM | 1 | 1 |
| | | WM-U | 3 | 2 |
| | | WM Deep | 3 | 2 |
| | iMOT(MRI) | Deep GM | 0 | 0 |

| | | | | |
|---|---|---|---|---|
| | | WM-U | 2 | 2 |
| | | WM Deep | 2 | 2 |
| | IPFC | Deep GM | 1 | 1 |
| | | WM-U | 3 | 2 |
| | | WM Deep | 3 | 2 |
| | MOT | Deep GM | 0.5 | 0 |
| | | WM-U | 2 | 2 |
| | | WM Deep | 1 | 2 |
| | SENS | Deep GM | 3 | 2 |
| | | WM-U | 3 | 2 |
| | | WM Deep | 3 | 2 |
| 6 (PSP) | ATP | Deep GM | 1 | 1 |
| | | WM-U | 3 | 3 |
| | | WM Deep | 3 | 3 |
| | OFC | Deep GM | 2 | 1 |
| | | WM-U | 3 | 3 |
| | | WM Deep | 3 | 3 |
| | IPFC | Deep GM | 2 | 2 |
| | | WM-U | 3 | 3 |
| | | WM Deep | 3 | 3 |
| | MOT | Deep GM | 3 | 2 |
| | | WM-U | 3 | 3 |
| | | WM Deep | 3 | 3 |
| | SENS | Deep GM | 3 | 3 |
| | | WM-U | 3 | 3 |
| | | WM Deep | 3 | 3 |
| 7(PiD) | OFC | Deep GM | 0 | 0 |
| | | WM-U | 1 | 1 |
| | | WM Deep | 1 | 1 |
| | DLPFC(MRI) | Deep GM | 0 | 0 |
| | | WM-U | 2 | 1 |
| | | WM Deep | 1 | 0.5 |
| | IPFC | Deep GM | 0.5 | 0.5 |
| | | WM-U | 1 | 1 |
| | | WM Deep | 1 | 1 |
| | MOT | Deep GM | 3 | 1 |
| | | WM-U | 2 | 2 |
| | | WM Deep | 2 | 2 |
| | SENS | Deep GM | 1 | 1 |
| | | WM-U | 3 | 2 |
| | | WM Deep | 3 | 2 |

Table depicts ordinal ratings of white matter in each patient sample region on standard ordinal rating scale (0=severe myelin degeneration with no discernable fibers, 0.5= severe myelin degeneration with rare isolated fibers, 1=moderate myelin degeneration, 2= mild myelin degeneration, 3=healthy appearing myelin without degeneration). WM LFB= white matter detected by luxol-fast blue myelin stain, WM Fe= white matter detected by Perl's stain for iron with DAB enhancement, HC= healthy control, AD= Alzheimer's disease, TDPA= FTLD-TDP subtype A, TDPC= FTLD-TDP subtype C, GGT= FTLD-Tau with globular glial tauopathy, PSP= FTLD-Tau with progressive supranuclear palsy, PiD= FTLD-Tau with Pick's disease, ATP= anterior temporal pole, OFC= orbitofrontal cortex, IPFC= inferior prefrontal cortex, MOT= primary motor cortex, SENS= primary sensory cortex, lOFC (MRI)= MRI guided sample from lateral OFC, iMOT (MRI)= MRI guided inferior primary motor cortex, DLPFC (MRI)= MRI guided sample from dorsolateral frontal cortex. deep GM= WM in deep cortical layers (stria Gennari or Baillarger), WM-U= juxtacortical white matter enriched for U-fibers, WM deep= relative deep WM.

### Supplementary Table 3. T2*w MRI contrast in MRI-guided sampling regions

| Region | Feature | Pt#1 HC | Pt#2 AD | Pt#33 TDPC | Pt#4 TDPA | Pt#5 Tau GGT | Pt#6 Tau PSP | Pt#7 Tau PiD |
|---|---|---|---|---|---|---|---|---|
| ATC | WM Subj./Overall | - | -1 | 0 | 0 | 0 | 0 | - |
| | WM UF/Subj. | - | -1 | -1 | -2 | 0 | 0 | - |
| | Upper/Lower Layers | - | 0 | 0 | 0 | 0 | 0 | - |
| | BoB/GM | - | -1 | -1 | 0 | -1 | -2 | - |
| | GM/Subj. WM | - | +2 | +2 | +2 | +2 | +2 | - |
| | Atyp. Features | - | DS | DS, UB, WS | DS, MB, WS | - | DS++ | - |
| OFC | WM Subj./Overall | 0 | 0 | 0 | 0 | 0 | 0 | +2 |
| | WM UF/Subj. | -2 | -1 | -1 | -2 | -1 | 0 | -2 |
| | Upper/Lower Layers | 0 | 0 | +1 | 0 | 0 | 0 | 0 |
| | BoB/GM | 0 | 0 | 0 | -1 | 0 | -1 | 0 |
| | GM/Subj. WM | +2 | +2 | +2 | +1 | +2 | +2 | 0 |
| | Atyp. Features | | | | DS | | | |
| IPFC | WM Subj./Overall | 0 | 0 | 0 | 0 | 0 | 0 | +1 |
| | WM UF/Subj. | -2 | 0 | -1 | -1 | 0 | 0 | -1 |
| | Upper/Lower Layers | +2 | +1 | +2 | +2 | +2 | 0 | 0 |
| | BoB/GM | -1 | -1 | -2 | -1 | -1 | -1 | -1 |
| | GM/Subj. WM | +2 | +2 | +2 | +2 | +2 | +2 | +2 |
| | Atyp. Features | | | | | | DS | |
| MOT | WM Subj./Overall | 0 | 0 | 0 | 0 | -2 | -1 | 0 |
| | WM UF/Subj. | -1 | -1 | -1 | 0 | 0 | 0 | 0 |
| | Upper/Lower Layers | +2 | +1 | +2 | +2 | +2 | +2 | +2 |
| | BoB/GM | -2 | -1 | -2 | -1 | a | a | -1 |
| | GM/Subj. WM | +2 | +2 | +2 | +2 | +2 | +2 | +2 |
| | Atyp. Features | | | | | MB++, WS | MB, WS | |
| SENS | WM Subj./Overall | 0 | 0 | 0 | 0 | 0 | 0 | 0 |
| | WM UF/Subj. | -1 | -1 | -1 | 0 | -1 | 0 | 0 |
| | Upper/Lower Layers | +2 | +1 | +2 | +2 | +1 | +2 | +2 |
| | BoB/GM | -2 | -1 | -2 | -1 | -2 | -2 | -1 |
| | GM/Subj. WM | +2 | +2 | +2 | +2 | +2 | +2 | +2 |
| | Atyp. Features | | | | | | | |

HC= Healthy Control; AD= Alzheimer's disease; TDPC= Frontotemporal Lobar Degeneration (FTLD) TDP-43 type C; TDPA= FTLD TDP-43 type A; Tau GGT= FTLD Tauopathy Globular Glial Tauopathy; Tau PSP= FTLD Tauopathy Progressive Supranuclear Palsy; Tau PiD= FTLD Tauopathy Pick's Disease; ATP= Anterior temporal pole; OFC= orbitofrontal cortex; IPFC= inferior prefrontal cortex; MOT= primary motor cortex; SENS= primary sensory cortex; DS = diffuse speckling, MB = mid-cortical hypointense band, WS = striated white matter; a = BoB was completely occluded by mid-cortical hypointense band through entire gyrus.

### Supplementary Table 4. T2*w MRI contrast in MRI-guided sampling regions

| Feature | Pt#4<br>TDPA<br>LOFC | Pt#5<br>Tau GGT<br>IMOT | Pt#6<br>Tau PiD<br>ILPFC |
|---|---|---|---|
| WM Subj./Overall | 0 | -1 | +2 |
| WM UF/Subj. | -2 | 0 | -1 |
| Upper/Lower Layers | +1 | +1 | 0 |
| BoB/GM | -2 | a | 0 |
| GM/Subj. WM | +2 | +2 | +1 |
| Atyp. Features | UB | MB++, WS | MB |

TDPA= FTLD TDP-43 type A; Tau GGT= FTLD Tauopathy Globular Glial Tauopathy; Tau PiD= FTLD Tauopathy Pick's Disease; LOFC = lateral orbitofrontal cortex; IMOT= inferior motor cortex; ILPFC = inferior lateral prefrontal cortex; UB = upper-cortical hypointense band, WS = striated white matter; a = BoB was completely occluded by mid-cortical hypointense band through entire gyrus

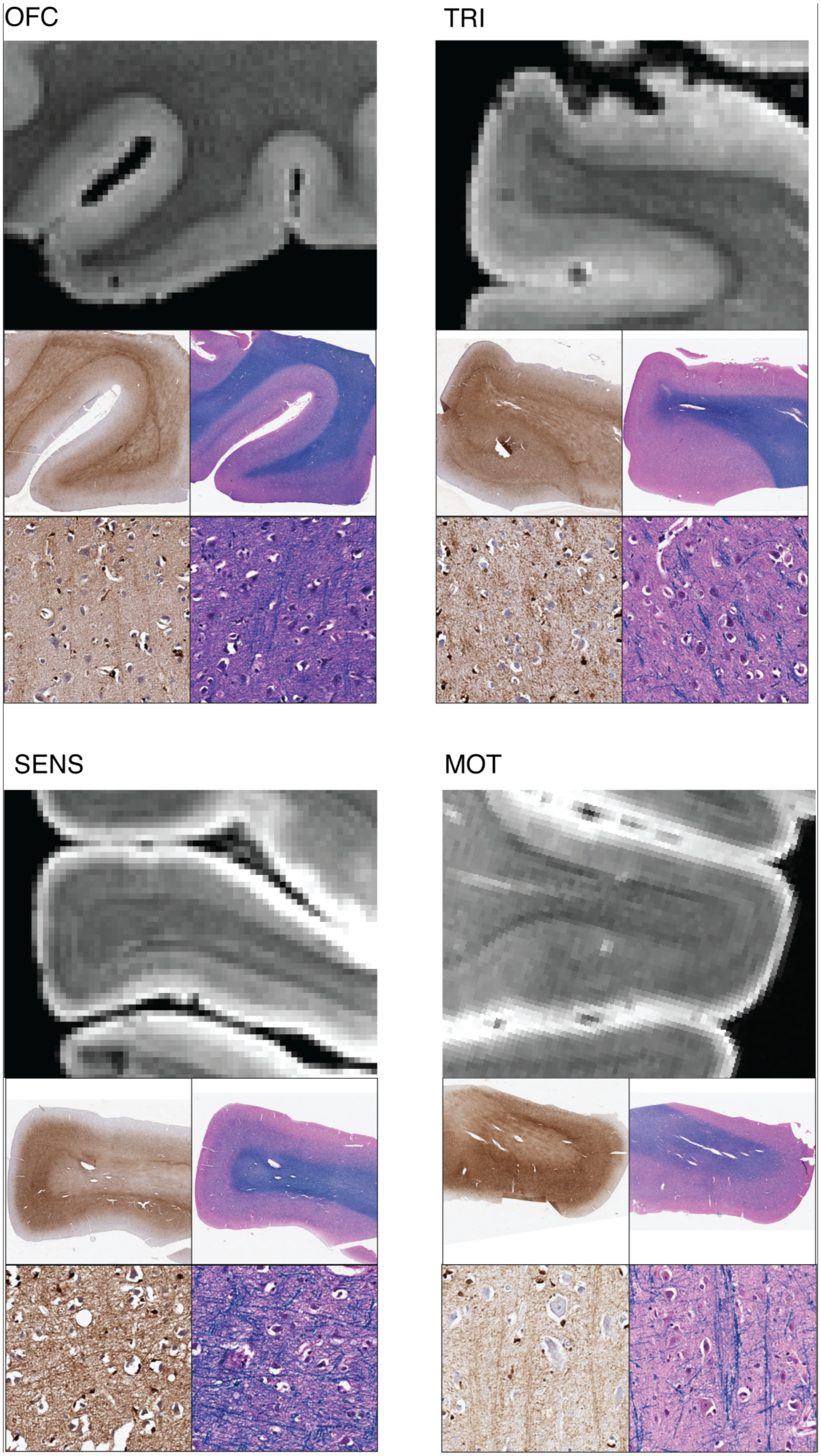
OFC
TRI
SENS
MOT

**Supplementary Figure 1. Myleoarchitecture of cortex in healthy control hemisphere.** In four representative regions we show T2*w MRI, co-located serial sections stained with iron and LFB (myelin) stain, and high-mag zoomed images of iron and LFB (myelin) stain from the deep cortical GM these same slices. Bands of Baillarger are visible as hypointense tangential stripes in the middle layers of the cortex, while radial fibers produce a moderate hypointensity of lower-layer cortical GM relative to more hyperintense upper layers. The myeloarchitecture varies from region to region, producing specific localized laminar pattern of T2*w image contrast.

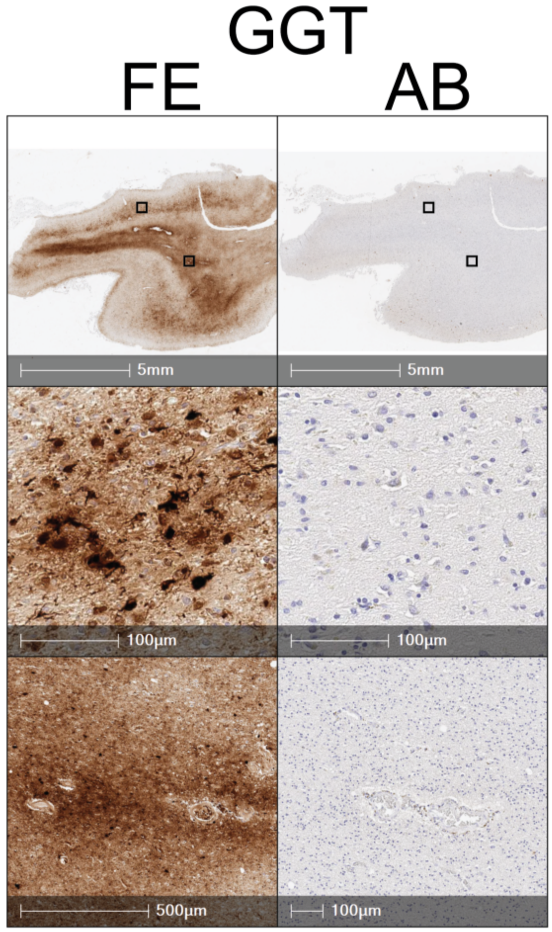


**Supplementary Figure 2. Pathological T2* signal in FTLD-Tau cases does not correspond to co-morbid amyloid plaque pathology.** Photomicrographs depict semi adjacent sections from 4R tauopathy (GGT, patient #5; Thal phase A2 CERAD C0) and progressive supranuclear palsy 4R tauopathy (PSP, patient #6; Thal phase A2 CERAD C3) stained for iron (FE) and amyloid-beta (AB). Top row of histology sections depicts 1x magnification and bottom rows depicts 32 x high magnification views from pathological areas (boxes). Pathological iron in mid to deep layers (arrows) corresponds to clusters of iron-rich microglia and not to the mild amounts of amyloid plaque pathology that was absent in these areas, suggesting 7T patterns observed in these cases is driven by pathophysiology of 4R tauopathy and not mild plaque co-pathology.

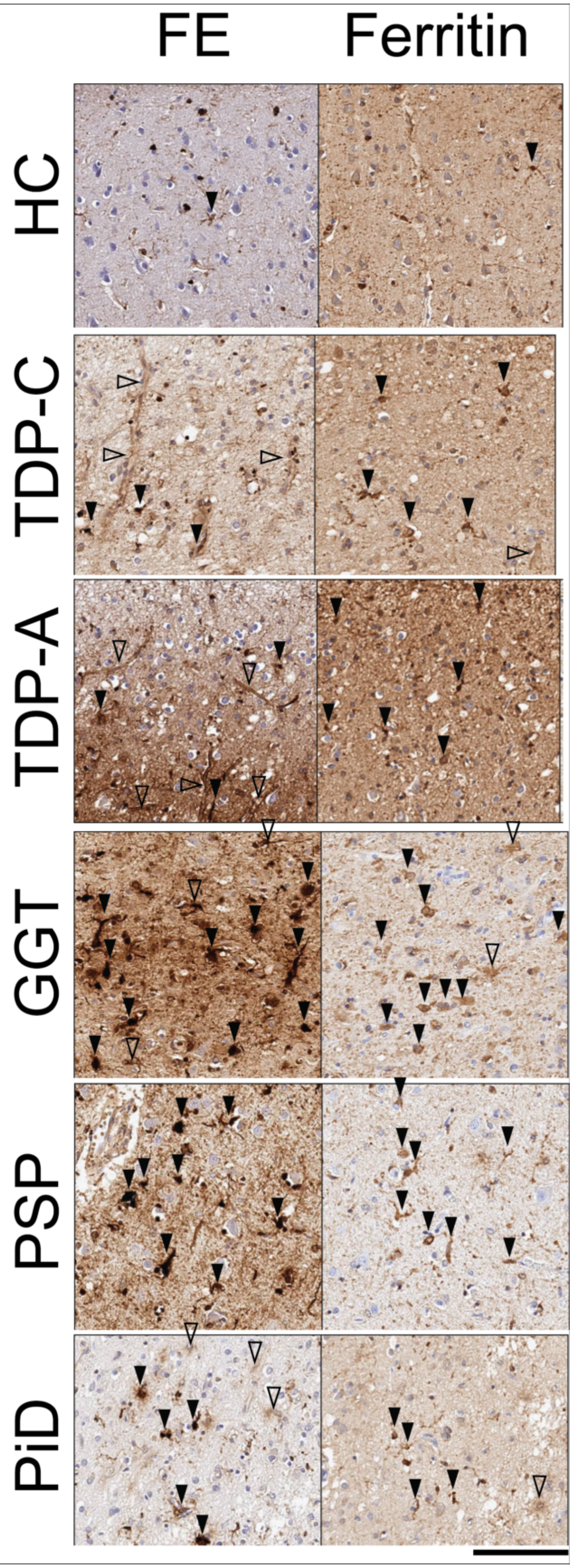
FE
Ferritin
HC
TDP-C
TDP-A
GGT
PSP
PiD

**Supplementary Figure 3. Comparison of DAB-enhanced Perl's iron staining and ferritin immunostaining in FTLD samples.** Photomicrographs depict iron-stained sections from regions of abnormal T2* signal ex vivo in FTLD and semi-adjacent sections immunostained for ferritin. The healthy control (HC) sample had iron reactivity largely limited to oligodendrocytes and healthy white matter with only rare iron- or ferritin-positive microglia. In FTLD samples the iron-rich microglia (black arrowheads) were similarly detected in ferritin immunostaining, while iron rich processes corresponding to astrocytes (open arrowheads) were minimally detected with ferritin immunostaining, likely due to known minimal ferritin content in astrocytes compared to microglia and oligodendrocytes.
